# ^13^C metabolite tracing reveals glutamine and acetate as critical in vivo fuels for CD8^+^ T cells

**DOI:** 10.1101/2023.06.09.544407

**Authors:** Eric H. Ma, Michael S. Dahabieh, Lisa M. DeCamp, Irem Kaymak, Susan M. Kitchen-Goosen, Dominic G. Roy, Mark J. Verway, Radia M. Johnson, Bozena Samborska, Catherine A. Scullion, Mya Steadman, Matthew Vos, Thomas P. Roddy, Connie M. Krawczyk, Kelsey S. Williams, Ryan D. Sheldon, Russell G. Jones

**Author notes:** Corresponding author: Russell G. Jones.

## Abstract

Infusion of 13C-labeled metabolites provides a gold-standard for understanding the metabolic processes used by T cells during immune responses *in vivo*. Through infusion of 13C-labeled metabolites (glucose, glutamine, acetate) in *Listeria monocytogenes* (*Lm*)-infected mice, we demonstrate that CD8+ T effector (Teff) cells utilize metabolites for specific pathways during specific phases of activation. Highly proliferative early Teff cells *in vivo* shunt glucose primarily towards nucleotide synthesis and leverage glutamine anaplerosis in the tricarboxylic acid (TCA) cycle to support ATP and *de novo* pyrimidine synthesis. Additionally, early Teff cells rely on glutamic-oxaloacetic transaminase 1 (Got1)—which regulates *de novo* aspartate synthesis—for effector cell expansion *in vivo*. Importantly, Teff cells change fuel preference over the course of infection, switching from glutamine-to acetate-dependent TCA cycle metabolism late in infection. This study provides insights into the dynamics of Teff metabolism, illuminating distinct pathways of fuel consumption associated with Teff cell function *in vivo*.

**Teaser:** Interrogating dynamics of fuel utilization by CD8^+^ T cells *in vivo* reveals new metabolic checkpoints for immune function *in vivo*.

## Introduction

The transition of quiescent naïve CD8^+^ T cells (Tn) towards an activated cell state is marked by pronounced transcriptional and metabolic changes that support rapid cell division and acquisition of effector functions (*1, 2*). This transition−often referred to as metabolic reprogramming−is underpinned by preferential uptake and processing of nutrients to support cellular metabolism (*3–5*). Stable isotope labeling (SIL) techniques using ^13^C-labeled substrates have enabled detailed mapping of intracellular nutrient utilization by immune cells (*6, 7*). Glucose is the most widely documented nutrient fueling T cell activation. Glycolytic breakdown and oxidation of glucose fuels energy (ATP) production (*8*), while secondary metabolism of glycolytic intermediates are used for post-translational modifications (i.e. histone acetylation) (*9*), biosynthetic programs that support proliferation (i.e. serine, nucleotide synthesis) (*10, 11*), and cell signaling (*12–14*). Targeting key metabolic nodes such as glucose uptake (*15, 16*), lactate metabolism (*17, 18*), one-carbon metabolism (*11, 14*), and mechanistic target of rapamycin complex 1 (mTORC1) signaling (*19–21*) modulate T cell responses, highlighting the importance of metabolic programs to T cell function.

Deciphering the cellular fates of fuels such as glucose has been modeled largely by applying SIL approaches to *in vitro* culture systems, which are limited by the use of supraphysiologic nutrient levels and limited metabolic diversity that fail to recapitulate physiologic environments *in vivo* (*18, 22–24*). Environmental context is an important determinant of cellular metabolic dependencies. Culturing T cells in medium that more closely models circulating metabolite levels in serum increases viability and effector cytokine (i.e., IFN-γ, TNF-α) production by human and mouse T cells, respectively (*18, 25*). Moreover, many substrates used to fuel oxidative metabolism in vitro are not prominently used in vivo. We previously reported that CD8^+^ Teff cells display distinct patterns of glucose utilization in vivo, using glucose as a substrate for biosynthesis (i.e., nucleotide, nucleotide sugar, and serine metabolism) but less for oxidative metabolism and lactate production (*10*). In contrast, lung tumors prominently oxidize glutamine for ATP production in vitro, but display minimal dependence on glutamine for tricarboxylic acid (TCA) cycle metabolism in vivo (*26*). Glutamine functions as a key anaplerotic substrate that replenishes TCA cycle intermediates in many cell types grown *in vitro*, including T cells (*27, 28*). However, how glutamine and other fuels are used by T cells *in vivo* has remained undefined.

Here we have used infusions of various ^13^C-labeled substrates (^13^C-glucose, glutamine, and acetate) to study nutrient utilization by CD8^+^ T cells *in vivo* over the course of *Listeria monocytogenes* (*Lm*) infection. We report that CD8^+^ T cells−in contrast to lung tumors−prominently use glutamine as an oxidizable fuel source *in vivo*. We find that glutamine anaplerosis into the TCA cycle and aspartate synthesis (via the enzyme glutamic-oxaloacetic transaminase (Got1)) are instrumental for CD8^+^ T cell ATP production and cell proliferation, respectively. However, mature CD8^+^ Teff cells at the peak of expansion reduce their reliance on glucose and glutamine as fuel sources, switching to acetate to fuel oxidative metabolism. These data highlight a critical role for glutaminolysis for early CD8^+^ T cell expansion and reveal plasticity in T cell fuel choice over the course of infection *in vivo*.

## Results

### Glutamine is the major TCA cycle fuel for CD8^+^ T cells *in vivo*

To study glutamine utilization by Teff cells *in vivo*, we examined the uptake and processing of ^13^C-labeled nutrients using stable isotope labeling (SIL) in mice infected with the Gram-positive bacterium *Listeria monocytogenes* (*Lm*). CD8^+^ OT-I T cells expressing a transgenic TCR specific for ovalbumin (OVA) and the congenic marker CD90.1 (Thy1.1) were adoptively transferred into naïve hosts, followed by infection with *Lm* expressing OVA (*Lm*OVA) (**Fig. 1a**). Mice were infused for ∼2-3 hours with either U-[^13^C]glucose or U-[^13^C]glutamine 3 days post infection (dpi), and naïve (Tn) or antigen-specific Teff cells isolated using magnetic beads following established protocols for rapid cell isolation (*10, 29*) (**Fig. 1a**). For the purposes of this study, “*in vitro*” denotes CD8^+^ OT-I T cells activated *in vitro* using standardized cell culture conditions (i.e., stimulation with OVA peptide and expansion with IL-2). “*In vivo*” indicates CD8^+^ OT-I T cells isolated from *Lm*OVA-infected mice following ^13^C-metabolite infusion, while cells analyzed “*ex vivo*” represents CD8^+^ OT-I T cells isolated from *Lm*OVA-infected mice subjected to short-term culture (< 4h) with medium containing ^13^C-labeled metabolites (*6*).

**Fig. 1.**
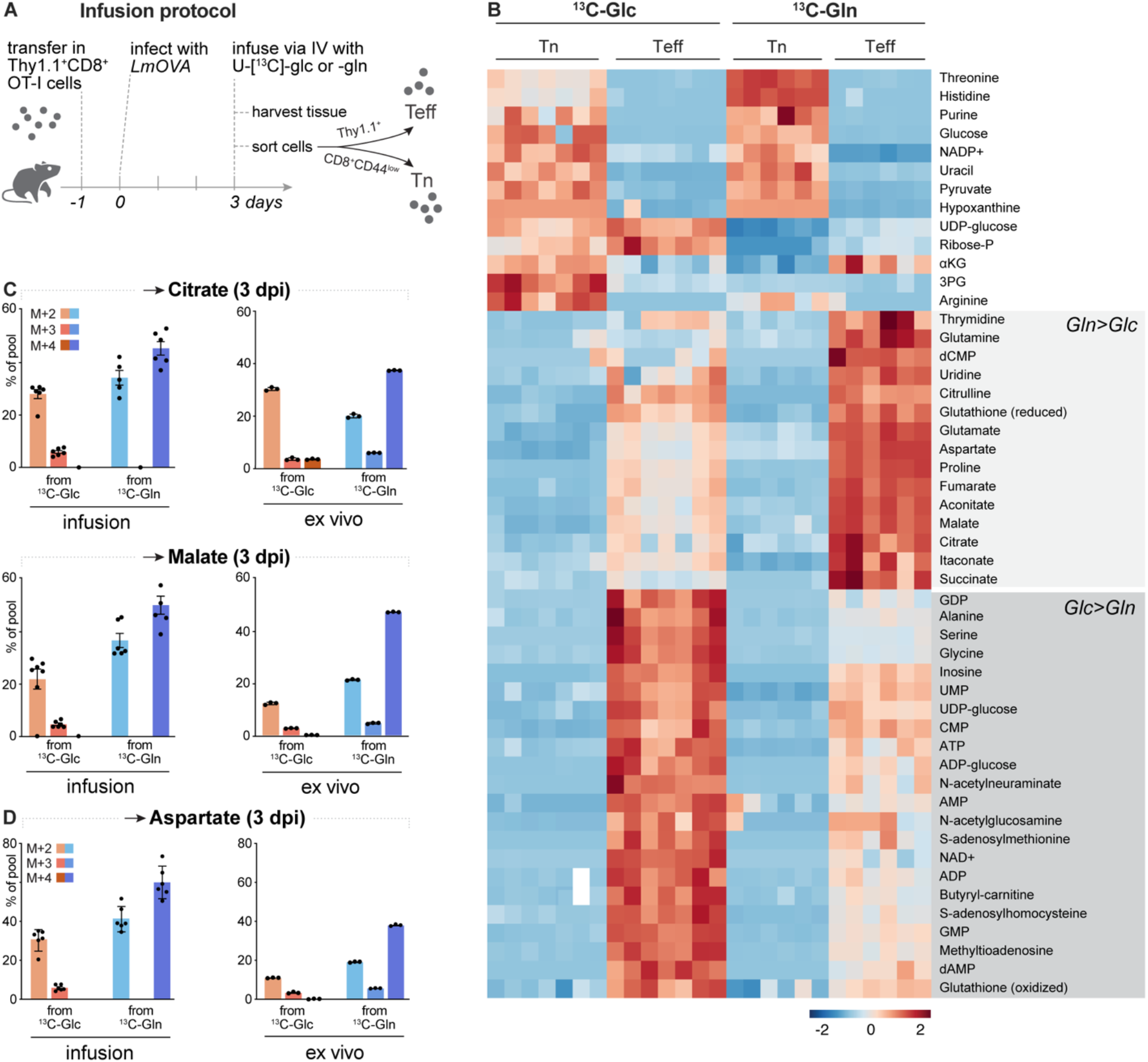
Glutamine is the major TCA cycle fuel for CD8^+^ T cells in vivo. **(A)** Infusion protocol schematic. One day prior to infection (day -1), Thy1.1^+^ CD8^+^ OT-I^+^ T (Teff) cells are transferred into sex-matched congenic Thy1.2^+^ recipients followed by infection with *LmOVA* the following day (day 0). On day 3, mice are anesthetized and infused with U-[^13^C]Glc or U-[^13^C]Gln. Homogenized spleens were split equally prior to Thy1.1^+^ (Teff) or naïve CD8^+^/CD44^low^ (Tn) cell selection. **(B)** Heat map depicting % labelling of intracellular metabolites isolated from Tn and Teff cells following ^13^C-Glc or ^13^C-Gln infusion. Regions of the heatmap with increased ^13^C enrichment from U-[^13^C]Gln (Gln>Glc) versus U-[^13^C]Glc (Glc>Gln) are indicated by light and dark grey shading, respectively (z-score range from -4 to 4). **(C-D)** Percent labeling from ^13^C-Glc or ^13^C-Gln (% of total pool) in intracellular **(C)** citrate (top) and malate (bottom) and **(D)** aspartate in CD8^+^ Teff cells isolated from *Lm*-OVA-infected mice (3 dpi) following 2h infusion (left) or cultured for 2h with ^13^C-Glc or ^13^C-Gln *ex vivo*. M+2, M+3, and M+4 isotopologues are shown. Data represent the mean ± SEM for biological replicates (n=3-6).

We first directly compared the use of glucose versus glutamine by CD8^+^ Teff cells responding to *Lm*OVA infection *in vivo*. Anesthetized mice were given a bolus of U- [^13^C]glutamine (0.3mg/gBW) followed by a continuous infusion (6×10^-3^ mg/min/gBW) for 150 min prior to isolation of CD8^+^ Thy1.1^+^ Teff or naïve T (Tn) cells from the spleens of infected animals **(Fig. 1a)**. Infusion of U-[^13^C]glutamine with this protocol allowed us to achieve ^13^C_5_- glutamine levels of ∼40% in the blood and liver, and ∼30% in the spleen **(Fig. S1a)**. ^13^C labeling of tricarboxylic acid (TCA) cycle intermediates (αKG, fumarate, succinate) and TCA cycle-derived amino acids (glutamate, aspartate) was also observed in the blood, liver, and spleen of *Lm*OVA-infected mice infused with U-[^13^C]glutamine **(Figure S1a)**.

Combining previously published work investigating the use of U-[^13^C]glucose infusion in CD8^+^ OT-I T cells responding to *Lm*OVA infection (*10*) with the infusion of U-[^13^C]glutamine, we identified metabolites in Tn or Teff cells with relative fractional enrichment from ^13^C-glucose and/or ^13^C-glutamine at 3 dpi **(Fig. 1b)**. The majority of central carbon metabolites contained heavy carbon (^13^C) from both glucose and glutamine; however, we observed distinct segregation of metabolites into groups that were preferentially labeled from either carbon source. Labeling from U-[^13^C]glucose was highly enriched in glycolytic intermediates and glycolysis-derived metabolites (i.e., purine/pyrimidine nucleotides and amino acids like serine and alanine) **(Fig. 1b**). Of note, *in vivo*-labelled Teff cells displayed a high level of labeling from ^13^C-glutamine in glutathione and intermediates of the TCA cycle (**Fig. 1b**). We estimated the contribution of glucose or glutamine to the first turn of the TCA cycle by measuring isotopologue distribution in TCA cycle metabolites: M+2 for glucose and M+4 for glutamine (see graphic in **Fig. S1b**). Heavy isotope labeling with U-[^13^C]glucose showed a M+2 label of approximately 30% in citrate and ∼20% in malate (**Fig. 1c**). In contrast, we observed a higher contribution of U-[^13^C]glutamine to the TCA cycle *in vivo* compared to glucose, with ∼45% of citrate and malate containing M+4 labelled carbon (**Fig. 1c**).

*Ex vivo* tracing of *Lm*OVA-specific CD8^+^ Teff cells using ^13^C-glucose or ^13^C-glutamine revealed similar patterns of nutrient utilization to that observed with *in vivo* infusion, with ^13^C- glutamine contributing more heavy carbon to citrate and malate production than ^13^C-glucose (**Fig. 1c** and **S1c**). In addition, the TCA cycle-derived amino acid aspartate—a critical component of pyrimidine nucleotide synthesis—was similarly enriched in ^13^C carbon from glutamine over glucose (**Fig. 1d** and **S1d**). Thus, in contrast to many tumor models (*26, 30*), glutamine is a significant fuel for T cell oxidative metabolism *in vivo* and contributes more carbon than glucose to the TCA cycle.

### OXPHOS is fueled largely from glutamine in physiologically activated CD8^+^ T cells

Glutamine plays an important role in the mitochondria through its anaplerotic functions in the TCA cycle (*31, 32*). Given the increased use of glutamine for TCA cycle metabolism by CD8^+^ Teff cells *in vivo* (**Fig. 1**), we examined differences in mitochondrial mass and activity between in vitro-activated versus physiologically-activated (via *Lm*OVA infection) CD8^+^ T cells fluorescent dyes that quantify mitochondrial mass (MitoSpy) versus mitochondria membrane potential (TMRM), respectively. CD8^+^ OT-I T cells activated *in vitro* displayed a ∼2.5-fold increase in mitochondrial mass compared to OT-I T cells responding to *Lm*OVA at 3 dpi (**Fig. 2a**). Despite this, OT-I T cells responding to *Lm*OVA displayed higher TMRM staining, indicating an overall increase in mitochondrial membrane potential relative to *in vitro*-activated T cells (**Fig. 2b**). Overall, this analysis revealed that OT-I cells responding to pathogen infection *in vivo* displayed higher mitochondrial activity (>5-fold higher) on a per-cell basis than *in vitro*-activated CD8^+^ OT-I T cells (**Fig. 2c**). These data may reflect a greater reliance on TCA cycle metabolism or use of diverse fuels−including glutamine−for mitochondrial function by CD8^+^ T cells *in vivo.* Glutamine anaplerosis replenishes the TCA cycle intermediates, providing reducing equivalents to feed the electron transport chain, generating energy through oxidative phosphorylation (OXPHOS) in the mitochondria.

**Fig. 2.**
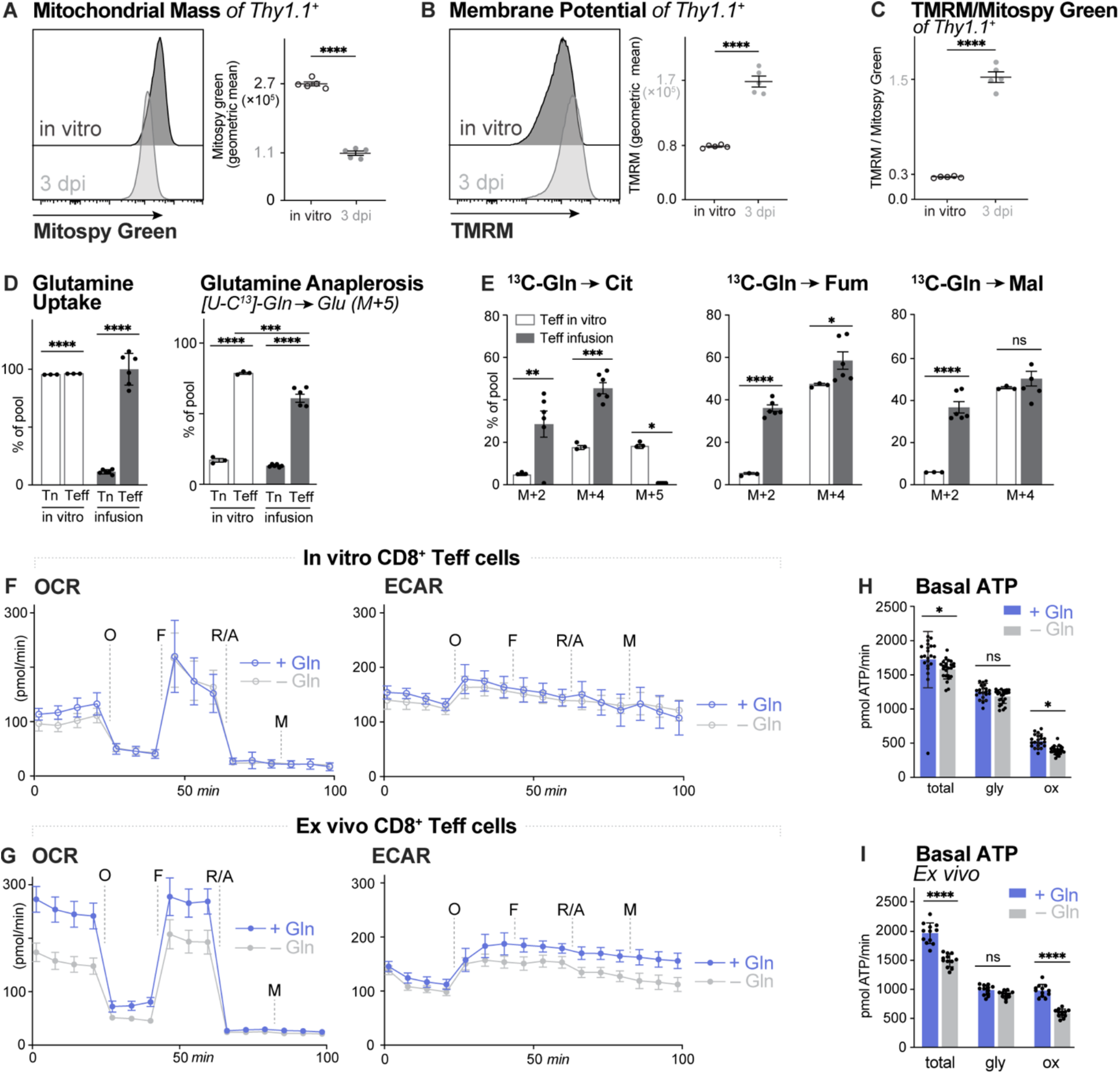
Glutamine fuels OXPHOS in physiologically activated CD8^+^ T cells (A-B) Histograms for Mitospy Green (**A**) and TMRM (**B**) fluorescence emission from CD8^+^ Teff cells following 3 days of *in vitro* activation or isolated from *Lm*-OVA-infected mice (3 dpi). Geometric mean fluorescence intensity (MFI) of Mitospy Green or TMRM staining between conditions are shown (mean ± SEM, n=5). **(C)** Ratio of TMRM/Mitosopy Green fluorescence for cells in (**A-B**). **(D)** (left) Glutamine (m+5) uptake within Tn or Teff cells upon *in vitro* culture with ^13^C-Gln or *in vivo* ^13^C-Gln infusion; plotted as % of M+5 intracellular glutamine pool normalized to external (ext.) M+5 glutamine in media (*in vitro*) and in serum (infusion); (right) Glutamine anaplerosis plot measuring glutamate (M+5) percent of intracellular pool normalized to M+5 glutamine levels in media (*in vitro*) and in serum (infusion). Data represent the mean ± SEM for biological replicates (n=3-6). **(E)** Measurements of ^13^C-Gln conversion into Citrate (Cit), Fumarate (Fum) and Malate (Mal) for *in vitro*-cultured Teff cells or Teff cells isolated from *Lm*-OVA- infected mice following infusion; M+N isotopologues are plotted as % of pool and normalized to external (ext.) M+5 glutamine in media (*in vitro*) or serum (infusion). Data represent the mean ± SEM for biological replicates (n=3-6). **(F-G)** Plots of oxygen consumption rates (OCR, left) and extracellular acidification rates (ECAR, right) for OT-I T cells **(F)** activated *in vitro* for 3 days or **(G)** isolated from *Lm*-OVA-infected mice (3 dpi). Plots in **(F)** and **(G)** are shown for T cells cultured in the presence (blue) or absence (gray) of glutamine. **(H)** Total, glycolytic (gly), and OXPHOS (Ox) contributions to basal ATP production (pmol ATP/min) for OT-I T cells activated *in vitro* **(H)** or isolated from *Lm*-OVA-infected mice (3 dpi) **(I)**. Data represent the mean ± SD (n=22-25).

We next investigated the dynamics of glutamine metabolism by T cells *in vivo* using SIL. At 3 dpi, CD8^+^ OT-I T cells actively take up extracellular glutamine, with ∼100% of intracellular glutamine fully labeled (M+5) when normalized to circulating levels of U-[^13^C]glutamine in serum (**Fig. 2d**). Glutamine enters the TCA cycle through conversion to glutamate and subsequently, conversion of glutamate to α-ketoglutarate by glutamate dehydrogenase (**Fig. S1b**). Approximately 60% of the intracellular glutamate pool in *Lm*OVA-responding Teff cells was derived from U-[^13^C]glutamine, showing similar patterns to *in vitro-*activated CD8^+^ OT-I T cells (**Fig. 2d**). Examination of MIDs for U-[^13^C]glutamine-derived metabolites revealed key differences in glutamine utilization by CD8^+^ T cells *in vivo*. CD8^+^ OT-I T cells from *Lm*OVA- infected mice displayed high abundance of M+2 and M+4 labeled TCA cycle intermediates citrate, fumarate, and malate, suggesting incorporation of U-[^13^C]glutamine carbon through one (M+4) and two (M+2) turns of the TCA cycle (**Fig. 2e**). This contrasted with *in vitro*-stimulated Teff cells, which displayed prominent M+4 but not M+2 labeling in these intermediates (**Fig. 2e**). Additionally, *in vitro*-stimulated Teff cells contained the presence of M+5 labeling in citrate−indicative of reductive carboxylation of α-ketoglutarate (*33, 34*)−that was not observed in Teff cells *in vivo* (**Fig. 2e**). Of note, *Lm*OVA-specific OT-I T cells cultured *ex vivo* with U-[^13^C]glutamine did not display reductive carboxylation (**Fig. S2a**), suggesting that reductive carboxylation is a feature of *in vitro*-stimulated T cells. Another difference is that Teff cells from U-[^13^C]glutamine-infused mice contained significantly higher levels of M+2 label in TCA cycle intermediates compared to *in vitro*-stimulated Teff cells **(Fig. 2e**), suggesting increased TCA cycling by activated CD8^+^ T cells *in vivo*. Increased flux through the TCA cycle would increase the production of reducing equivalents (i.e., NADH, FADH_2_) for electron transport and may explain the increased mitochondrial membrane potential of Teff cells proliferating *in vivo* (**Fig. 2b-c**).

Next, we used a Seahorse extracellular flux analyzer to measure the contribution of glutamine to the bioenergetics of CD8^+^ OT-I T cells responding to *Lm*OVA infection at 3 dpi. Basal OCR was significantly higher in OT-I T cells isolated from *Lm*OVA-infected mice (∼240 pmol/min) compared to *in vitro-*activated OT-I T cells (∼130 pmol/min) (**Fig. 2f-g**), corresponding with increased mitochondrial activity associated with elevated TMRM staining in OT-I T cells analyzed *ex vivo* **(Fig. 2b)**. We next analyzed the glutamine dependence of T cell bioenergetics by removing extracellular glutamine from the Seahorse medium. Dropping glutamine from the culture medium caused minimal changes to the ECAR (**Fig. 2f-g**) or ATP production rates from glycolysis (**Fig. 2h-i**) for both *in vitro*-activated and *Lm*OVA-responding OT-I T cells. However, physiologically activated OT-I T cells displayed high sensitivity to glutamine withdrawal, resulting in a ∼40% drop in OCR upon removal of glutamine (**Fig. 2g**). This analysis revealed a direct impact of glutamine availability on ATP production from OXPHOS−with no evidence of compensation from glycolytic ATP production−in OT-I T cells responding to pathogen infection *in* vivo (**Fig. 2i**). Together, these data indicate increased coupling of glutamine-dependent TCA cycle metabolism to oxidative ATP production in CD8^+^ Teff cells responding to pathogen infection *in vivo* compared to *in vitro*-stimulated T cells that do not depend on glutamine for oxidative ATP production.

### Teff cells use glutamine as a biosynthetic substrate

Glutamine oxidation not only replenishes TCA cycle intermediates, but glutamine carbon can also be used as a biosynthetic precursor, whereby carbons from glutamine are used to synthesize other amino acids like proline and aspartate **(Fig. 3a)** (*31*). We next assessed glutamine-dependent biosynthesis in CD8^+^ T cells *in vivo* by calculating the relative fractional enrichment of glutamine-derived metabolites relative to ^13^C-glutamine in the spleen (for *in vivo* infusions) or extracellular medium (*in vitro*-stimulated and *ex vivo* cultured cells). ^13^C-glutamine labeling into proline was not observed in Tn cells either *in vitro* or from *in vivo* infusions **(Fig. 3b)**; however, a substantial increase in ^13^C_5_-proline was observed in activated CD8^+^ T cells (both *in vitro* and from infusion experiments), with CD8^+^ Teff cells *in vivo* displaying higher relative enrichment of ^13^C-glutamine carbon into the proline pool **(Fig. 3b)**. Similar to proline, *de novo* biosynthesis of aspartate significantly increased upon activation in both *in vitro-* and *in vivo-*activated T cells **(Fig. 3c)**, with CD8^+^ Teff cells responding to *Lm*OVA infection *in vivo* displaying higher fractional enrichment of M+4 and M+2 isotopologues of aspartate from ^13^C-glutamine compared to *in vitro*-stimulated CD8^+^ T cells (**Fig. 3c**). We observed similar increases in M+2 and M+4 aspartate from ^13^C- glutamine in *Lm*OVA-responding infection CD8^+^ Teff cells analyzed *ex vivo* (**Fig. S3**), highlighting increased aspartate synthesis as a metabolic feature of CD8^+^ T cells *in vivo*.

**Fig. 3.**
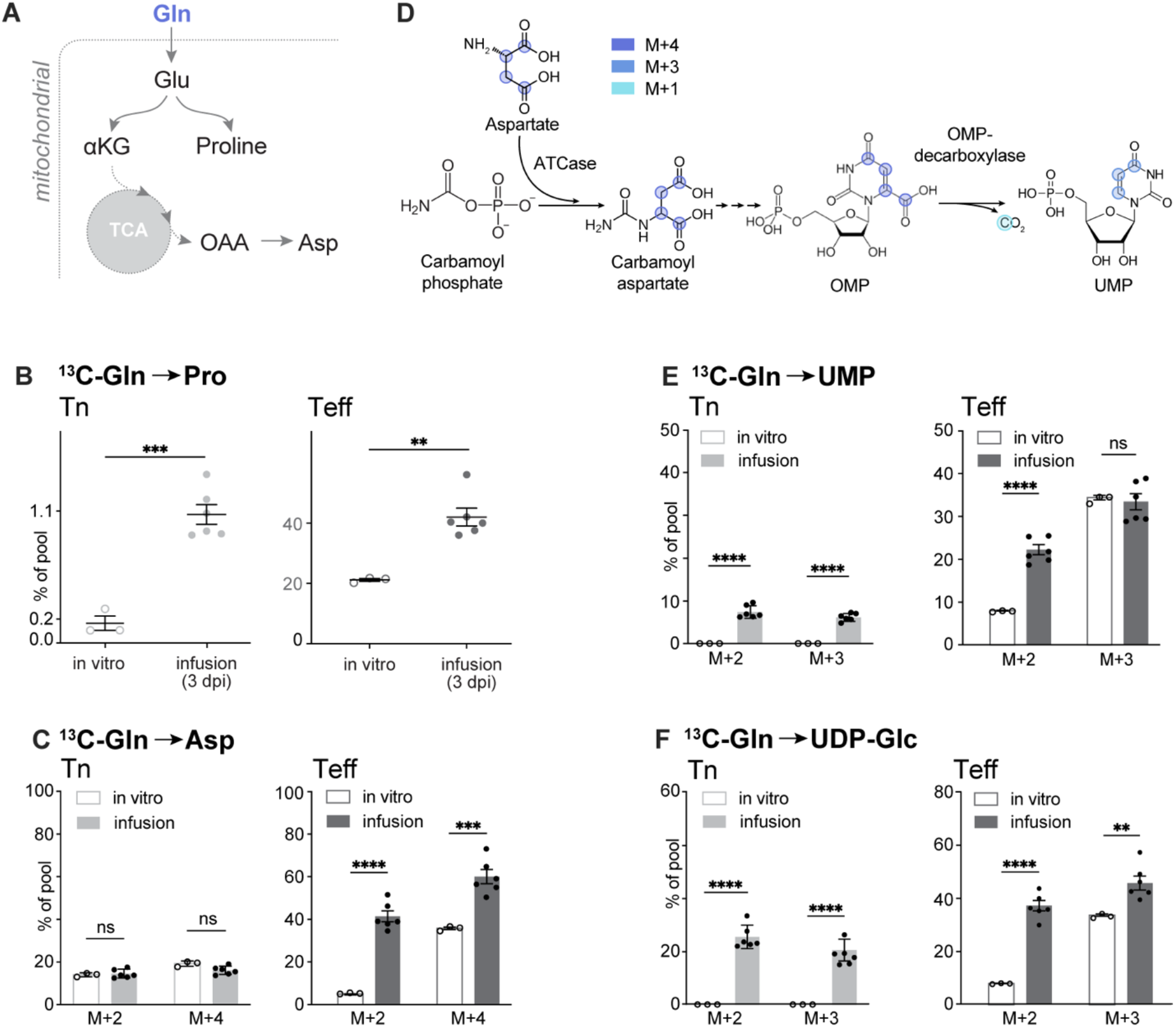
Glutamine is a substrate for amino acid and nucleotide biosynthesis in Teff cells. **(A)** Schematic depicting glutamine (Gln) carbon utilization for proline and aspartate (Asp) synthesis (Glu: Glutamate, αKG: α-ketoglutarate; OAA: oxaloacetate). **(B)** (left) ^13^C-Glutamine (Gln) conversion to proline by Tn cells (left) and Teff cells (right) following 2h *in vitro* culture with ^13^C-Gln or 2h *in vivo* ^13^C-Gln infusion. Total ^13^C enrichment in proline is plotted as the percentage of M+5 intracellular glutamine relative to external M+5 glutamine levels in media (*in vitro*) or serum (infusion). Data represent the mean ± SEM for biological replicates (n=3-6). **(C)** Conversion of ^13^C-Gln to aspartate for Tn and Teff cells treated as in **(B)**. Data are plotted as the percentage of M+2 and M+4 isotopologues normalized to M+5 glutamine in media *in vitro*) and in serum (infusion). **(D)** Schematic depicting the contribution of aspartate carbons to *de novo* pyrimidine (UMP) synthesis. ATCase, aspartate carbamoyltransferase; OMPdc, orotidine 5’- monophosphate decarboxylase. **(E-F)** ^13^C-Gln conversion into **(E)** UMP and **(F)** UDP-Glc for Tn and Teff cells displaying % of pools of M+2 and M+3. Data represent the mean ± SEM for biological replicates (n=3-6).

Aspartate is a non-essential amino acid important for proliferating cells as an intermediate in protein and nucleotide biosynthesis (*35*). The amine on aspartate is utilized for *de novo* purine nucleotides and the carbon backbone of aspartate is used for pyrimidine ring biosynthesis **(Fig. 3d)**. We analyzed the ^13^C labeling patterns in UMP to assess the contribution of ^13^C-glutamine-derived aspartate to pyrimidine nucleotide biosynthesis **(Figs. 3e-f)**. We observed prominent M+3 labeling of UMP from ^13^C-glutamine, with approximately 30% of UMP M+3 labeled in Teff cells both *in vitro* and from *in vivo* infusions **(Fig. 3e)**. In addition, we observed significant levels of M+2 labeled UMP in *Lm*OVA-responding Teff cells **(Fig. 3e)**, aligning with the labeling patterns observed in ^13^C-glutamine-derived aspartate *in vivo* **(Fig. 3c)**. The high prevalence of M+2 labeled aspartate and UMP in Teff cells *in vivo* likely corresponds to ^13^C-glutamine contributing to multiple rounds of TCA cycling. Tn cells, in contrast, displayed minimal ^13^C-glutamine labeling in UMP, likely due to the low nucleotide demand of non-proliferating cells **(Fig. 3e)**. ^13^C- glutamine-derived aspartate was also detected in nucleotide sugar UDP-glucose **(Fig. 3f**), which derives its nucleotide moiety from UMP. ^13^C-glutamine labeling in UDP-glucose followed that of UMP, with a larger proportion of M+2 UDP-glucose observed in Teff cells following *in vivo* infusion compared to *in vitro*-stimulated T cells **(Fig. 3f)**. Collectively, these data indicate a large proportion of glutamine carbon is used to drive TCA cycle-dependent biosynthetic reactions in CD8^+^ Teff cells *in vivo*.

### Got1 supports CD8^+^ Teff cell proliferation *in vivo*

The level of ^13^C-glutamine-derived aspartate generated by CD8^+^ Teff cells *in vivo* **(Fig. 3c)** suggests *de novo* synthesized aspartate may be important for Teff cell function. Aspartate is synthesized as part of the malate-aspartate shuttle, which functions to both shuttle metabolites and control the oxidation status of NAD^+^ between the cytosol and mitochondria **(Fig. 4a)**. The enzymes glutamic-oxaloacetic transaminase 1 and 2 (Got1 and Got2), which function in the cytosol and mitochondria, respectively, mediate the interconversion of oxaloacetate and glutamate to aspartate and α-ketoglutarate **(Fig. 4a)**. We utilized short hairpin RNA (shRNA)-mediated knockdown of Got1 in CD8^+^ OT-I T cells to test the metabolic and functional requirements for aspartate biosynthesis in CD8^+^ T cells. CD8^+^ T cells expressing Got1-targeting shRNAs displayed reduced Got1 protein expression **(Fig. 4b)**, as well as reduced overall abundance of ^13^C-glutamine-derived aspartate (**Fig. 4c**) including reduced levels of M+4 aspartate (**Fig. 4d**). Consistent with recent results shown for *Got1*^-/-^ T cells (*36*), silencing Got1 promoted a slight reduction in CD8^+^ T cell proliferation (**Fig. 4e**); however, the proliferation of Got1-knockdown—but not control—CD8^+^ T cells was selectively reduced when aspartate was removed from the cell culture medium **(Fig. 4e)**. These data suggest that T cells become auxotrophic for aspartate when Got1 gene expression is disrupted.

**Fig. 4.**
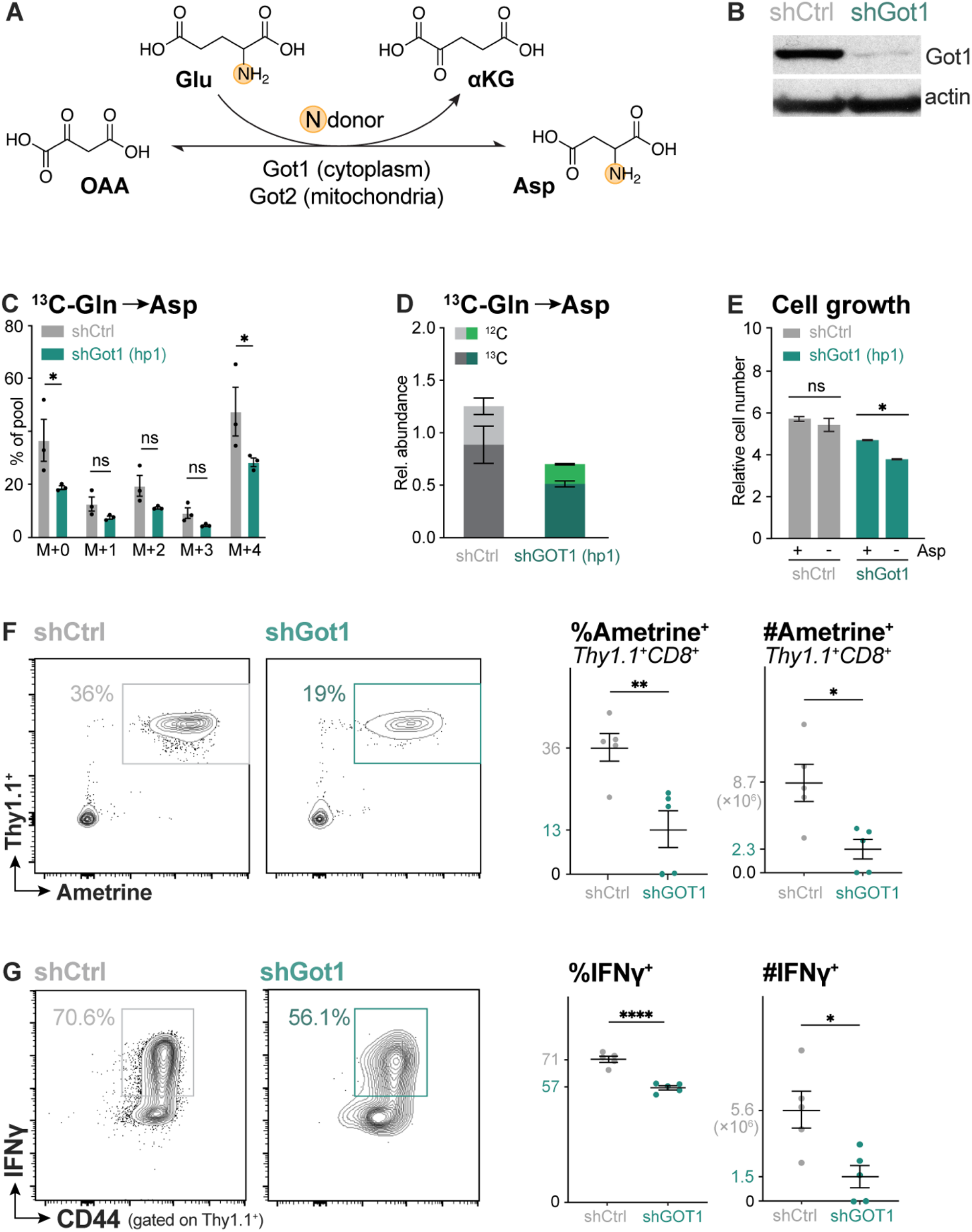
Got1 supports CD8^+^ T cell expansion *in vivo*. **(A)** Schematic of glutamic-oxaloacetic transaminase (Got) activity. Got1 (cytoplasm) and Got2 (mitochondria) catalyze the reversible transamination of oxaloacetate (OAA) to aspartate, using glutamic acid (Glu) as a nitrogen donor. **(B)** Immunoblot of Got1 and β-actin protein levels in CD8^+^ T cells expressing control (shCtrl) and *Got1*-targeting (shGot1) shRNAs. **(C)** Mass isotopologue distribution (MID) of ^13^C-Gln labeling into the intracellular aspartate pool in shCtrl- or shGot1-expressing CD8^+^ T cells. Data represent the mean ± SEM for biological replicates (n=3). **(D)** Total abundance of ^12^C- versus ^13^C-labeled intracellular aspartate in shCtrl- versus shGot1-expresssing CD8^+^ T cells expressing. T cells were cultured with ^13^C-Gln for 2h prior to metabolite extraction. Data represent the mean ± SEM for biological replicates. **(E)** Relative cell number for shCtrl- and shGot1-expressing T cells cultured for 2 days in medium containing (+) or lacking (-) aspartate (Asp). Data represent the mean ± SEM for biological replicates (n=3), with cell counts normalized to cell number at day 0. **(F-G)** Expansion and function of shGot1- expressing CD8^+^ T cells. Thy1.1^+^ OT-I T cells were transduced with vectors co-expressing Ametrine along with control or *Got1*-targeting shRNAs. Ametrine^+^ cells were adoptively transferred into Thy1.2^+^ hosts, followed by infection with *Lm*-OVA. **(F)** Thy1.1 versus Ametrine expression by CD8^+^ T cells at 6 dpi *Lm*-OVA infection expression. *Left,* representative flow cytometry plots for Thy1.1 and Ametrine expression. *Right*, percentage and number of shRNA- expressing (Ametrine^+^) *Lm*-OVA-specific T cells. **(G)** IFN-γ versus CD44 expression for shRNA- expressing OT-I T cells at 6 dpi. *Left,* representative flow cytometry plots for CD44 versus IFN-γ expression. *Right,* percentage and number of IFN-γ^+^ shRNA-expressing (Ametrine^+^) T cells. Data represent the mean ± SEM for biological replicates (n=5).

Next, we examined the role of Got1 in CD8^+^ T cell responses to *Lm*OVA infection *in vivo*. CD8^+^ OT-I T cells (Thy1.1^+^) transduced with control or Got1-targeting shRNAs were adoptively transferred into C57BL/6J (Thy1.2^+^) mice, followed by infection with *Lm*OVA one day later. At 7 dpi, we observed a marked reduction in both the percentage and number of antigen-specific (Thy1.1^+^) CD8^+^ T cells responding to *Lm*OVA infection upon Got1 silencing (**Fig. 4f**). Moreover, Got1 silencing altered the CD8^+^ T cell effector response, as evidenced by an overall lower percentage and number of IFN-γ-producing CD8^+^ T cells in the spleens of *Lm*OVA-infected animals (**Fig. 4g**). Together these data indicate a critical function for Got1 in mediating the expansion of CD8^+^ effector T cells *in vivo*, in a manner that cannot be compensated by Got2.

### Glutamine utilization by CD8^+^ T cells changes over the course of infection

One of the advantages of our *in vivo* infusion method is the ability to examine T cell metabolism *in situ* at specific times during the T cell response to infection (**Fig. 5a**). CD8^+^ T cells responding early to *Lm-*OVA infection (3 dpi) are predominantly early effector cells (EECs, KLRG1^lo^CD127^lo^), whereas Teff cells responding at the peak of the response (6-7 dpi) consist of a mix of EECs, short lived effector cells (SLECs, KLRG1^hi^CD127^lo^), and memory precursor effector cells (MPECs, KLRG1^lo^CD127^hi^) (*10*). We previously demonstrated that the shift from EECs to more differentiated CD8^+^ T cell subtypes (SLECs, MPECs) corresponds to a decrease in U-[^13^C]glucose utilization for both bioenergetic metabolism and anabolic growth pathways (*10*). We therefore examined whether CD8^+^ T cells display similar differences in glutamine usage *in vivo* using U-[^13^C]-glutamine infusions.

**Fig. 5.**
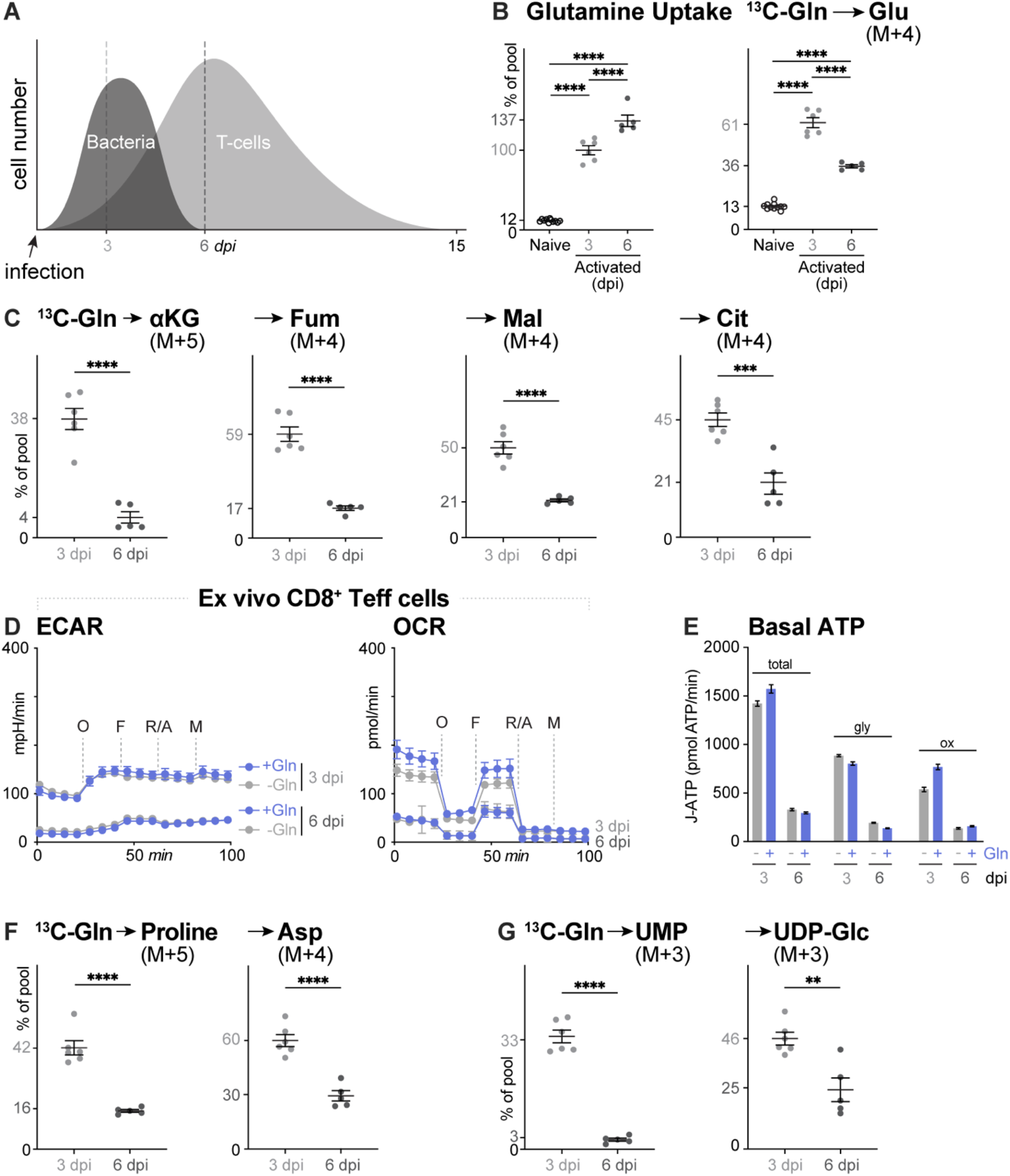
Glutamine utilization by CD8^+^ T cells changes over the course of infection. **(A)** Schematic of bacterial and CD8^+^ T cell numbers over the course of *Lm*-OVA infection. **(B-C)** Fractional enrichment of ^13^C-Gln into intracellular metabolite pools in naïve CD8^+^ T cells or OT- I Teff cells isolated from *Lm*-OVA infected mice at early (3 dpi) or peak (6 dpi) stages of the T cell response. Mice at each stage were infused with ^13^C-Gln for 2h prior to cell isolation and extraction of intracellular metabolites. Data represent the mean ± SEM (n=5-6). **(B)** ^13^C fractional enrichment into intracellular glutamine (M+5) and glutamate (M+5) pools. Data are normalized to serum ^13^C-Gln (M+5) levels. **(C)** ^13^C fractional enrichment into intracellular α-ketoglutarate (αKG), fumarate (fum), malate (mal) and citrate (cit) pools of Teff cells at 3 or 6 dpi. **(D-E)** Bioenergetics of CD8^+^ Teff cells at 3 and 6 dpi. **(D)** ECAR and OCR measurements for Teff cells isolated at 3 and 6 dpi and cultured in the presence (+, blue) or absence (-, grey) of glutamine. **(E)** Total, glycolytic (gly), and OXPHOS (Ox) contributions to basal ATP production (pmol ATP/min) in Teff cells isolated from *Lm*-OVA-infected mice (3 or 6 dpi). **(F-G)** ^13^C-glutamine (Gln) percent labelling into intracellular **(F)** proline (M+5) and aspartate (M+4) pools and **(G)** UMP (M+3) and UDP-Glc (M+3) pools in Teff cells at 3 and 6 dpi. Data are normalized to serum ^13^C-Gln (M+5) levels (mean ± SEM, n=5).

We calculated the relative fractional enrichment of glutamine-derived metabolites relative to the fractional enrichment of circulating U-[^13^C]glutamine present in the spleen, comparing first the uptake of U-[^13^C]glutamine by early (3 dpi) and late (6 dpi) CD8^+^ Teff cells and its subsequent conversation to ^13^C_5_-glutamate. U-[^13^C]glutamine was readily imported by both early and late Teff cells compared to Tn cells (relative fractional enrichment: early Teff cells, ∼1; late Teff cells, ∼1.3) **(Fig. 5b)**. A relative fractional enrichment >1 in late Teff cells suggests that glutamine is not only actively imported by late Teff cells, but that it is also either being stored in T cells or actively synthesized from ^13^C_5_-glutamate present in circulation during U-[^13^C]glutamine infusions **(Fig. S1a)**. Late Teff cells contain higher ^13^C-glutamine-derived M+5 glutamate (∼0.36) compared to Tn cells (∼0.13), indicating a higher glutamine-to-glutamate conversion, but this conversion was lower compared to early Teff cells (∼0.61) **(Fig. 5b)**. Surprisingly, while U-[^13^C]glutamine is actively imported by late Teff cells, we did not observe evidence of glutamine oxidation in the TCA cycle of late Teff cells, as noted by decreased fractional enrichment of U-[^13^C]glutamine-derived α-ketoglutarate (M+5), fumarate (M+4), malate (M+4), and citrate (M+4) at in late Teff cells compared to early Teff cells **(Fig. 5c)**. Similarly, we observed lower levels of M+2 labeled fumarate, malate, and citrate from U-[^13^C]glutamine in late versus early Teff cells, which suggests a decrease in TCA cycling in Teff cells at the peak response to infection **(Fig. S4a)**.

Comparing the bioenergetic profiles of *Lm*OVA-specific CD8^+^ Teff cells using the Seahorse extracellular flux analyzer, we found Teff cells at the peak response to infection (6 dpi) display significantly lower ECAR and OCR compared to early Teff cells (3 dpi) undergoing clonal expansion **(Fig. 5d)**. This corresponded to a roughly 6-fold reduction in ATP production capacity−from both glycolysis and OXPHOS−by CD8^+^ T cells at 6 dpi (**Fig. 5e**). These data align with the reduced utilization of both U-[^13^C]glucose (*10*) and U-[^13^C]glutamine (**Fig. 5c**) by Teff cells at the peak of the effector T cell response as determined by infusion of ^13^C-labeled metabolites into mice. Notably, the ATP production capacity was low despite the maintenance of robust effector function in Teff cells at 6 dpi (*10*). While the OCR of early Teff cells at 3 dpi was sensitive to glutamine withdrawal (**Fig. 2g**), both the OCR and ATP production from OXPHOS were unaffected by glutamine withdrawal in late Teff cells **(Figs. 5d-e)**.

Finally, we assessed the contribution of U-[^13^C]glutamine carbon to macromolecular biosynthesis in late Teff cells in U-[^13^C]glutamine-infused mice. Compared to early Teff cells at 3 dpi, late Teff cells at 6 dpi displayed reduced glutamine-dependent amino acid synthesis (i.e, proline and aspartate, **Fig. 5f**) and labeling into UMP and UDP-glucose (**Fig. 5g**). Together these data indicate that Teff cells reduce their reliance on glutamine for OXPHOS-dependent ATP production and TCA cycle-dependent biosynthesis by the peak of the effector response.

### Glutamine utilization by CD8^+^ T cells correlates with proliferative rate

To further explore the differences in glutamine utilization by CD8^+^ T cells over the course of infection, we conducted *ex vivo* analysis of antigen-specific CD8^+^ Teff cells at different stages during the response to *Lm*OVA infection. Using Ki67 as a measure of T cell proliferation, we observed similar proliferation rates between early Teff cells at 3 dpi and CD8^+^ T cells activated *in vitro* (**Fig. 6a**); however, we observed a dramatic 90-fold reduction in Ki67 staining in late Teff cells at 6 dpi compared to early Teff cells (**Fig. 6a**), indicating that Teff cells at the peak of the T cell response are a slowly proliferating population.

**Fig. 6.**
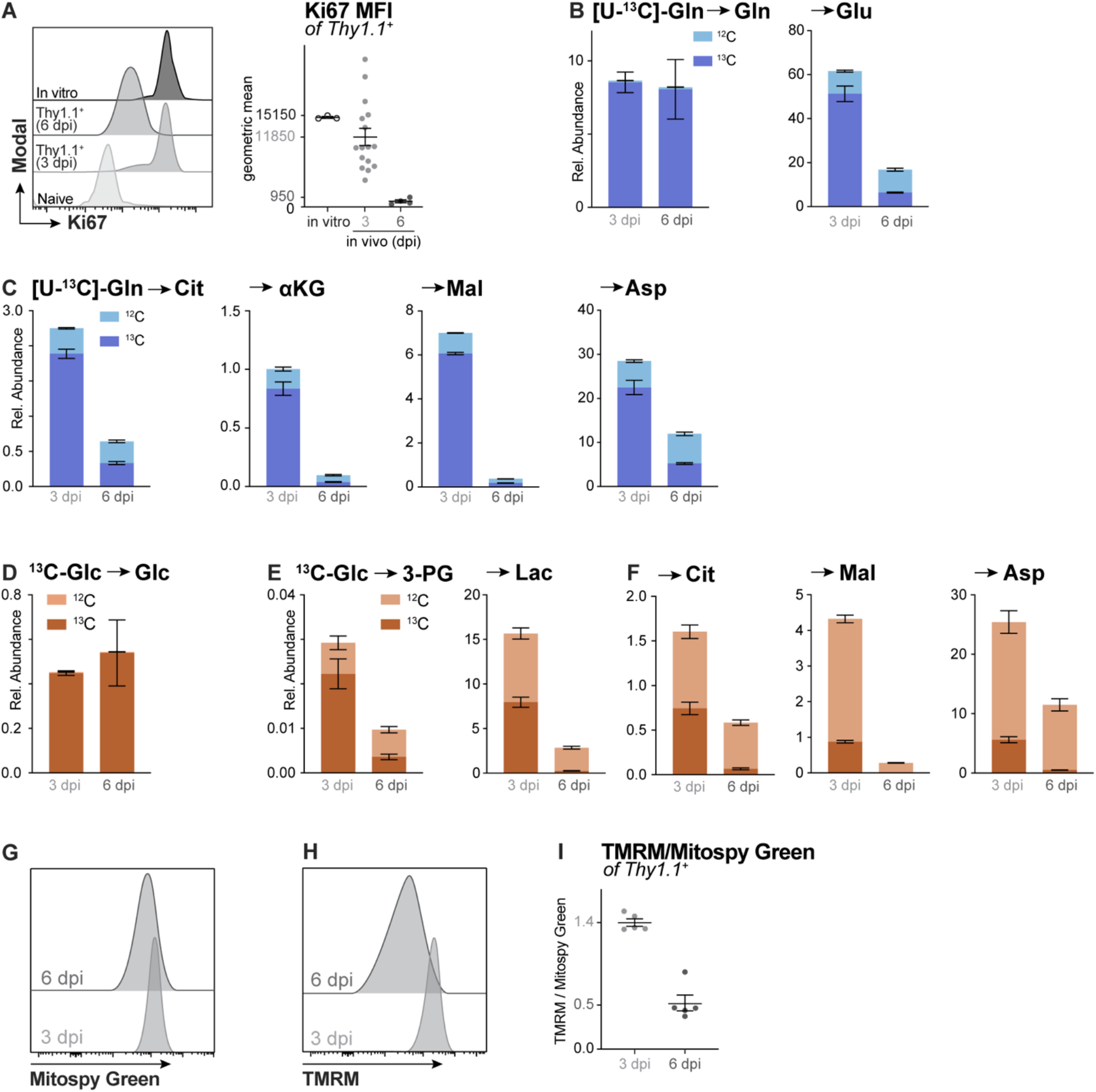
Glutamine utilization by CD8^+^ T cells correlates with proliferation rate. **(A)** Ki67 staining in CD8^+^ Teff cells. *Left,* Ki67 fluorescence emission (modal) histograms for resting (naïve) CD8^+^ T cells, *in vitro*-activated T cells (anti-CD3/CD28 stimulation for 3 days), or *in vivo*-activated Teff cells (isolated from *Lm*-OVA-infected mice 3 or 6 dpi). *Right,* geometric mean fluorescence intensity (MFI) for Ki67 staining. Data represent the mean ± SEM, n=3-16. **(B- C)** Relative abundance of ^12^C- and ^13^C-labelling in intracellular metabolite pools in CD8^+^ Teff cells isolated from *Lm*-OVA-infected mice that received [U-^13^C]Gln infusions at 3 or 6 dpi. **(B)** Abundance of ^12^C- versus ^13^C-labeled Gln (left) and Glu (right) in Teff cells. **(C)** Abundance of ^12^C- versus ^13^C-labeled citrate (Cit), α-ketoglutarate (αKG), malate (Mal), and aspartate (Asp). **(D-E)** Relative abundance of ^12^C- and ^13^C-labelling in intracellular metabolite pools in CD8^+^ Teff cells isolated from *Lm*-OVA-infected mice that received [U-^13^C]-Glc infusions at 3 or 6 dpi. **(D)** Abundance of ^12^C- versus ^13^C-labeled intracellular glucose (Glc) following infusion. **(E)** Abundance of ^12^C- versus ^13^C-labeled intracellular 3-phosphoglycerate (3-PG), lactate (Lac), citrate (Cit), malate (Mal), and aspartate (Asp). **(G-I)** Mitochondrial mass (Mitospy) and membrane potential (TMRM) of Teff cells isolated from *Lm*-OVA-infected mice at 3 or 6 dpi. Histograms of **(G)** Mitospy Green (mitochondrial mass) and **(H)** TMRM (membrane potential) fluorescence emission. **(I)** Ratio of TMRM/Mitosopy Green in Teff cells at 3 or 6 dpi. Data represent the mean ± SEM for biological replicates (n=5).

We next examined ^13^C-glutamine and ^13^C-glucose utilization in these *ex vivo* sorted Teff cell populations from different timepoints of infection. Due to the bead sorting procedure following *in vivo* infusion, we are only able to quantify the fractional enrichment of each metabolite, as the metabolite pool sizes is an unreliable readout due to loss of intracellular metabolites to exchanges with the sorting medium (*10, 29*). Isolated Teff cells were cultured for 2 hours in U-[^13^C]glutamine or U-[^13^C]glucose medium *in vitro*. Both early (3 dpi) and late (6 dpi) Teff cells displayed comparable levels of glutamine uptake, yet glutamine-to-glutamate conversion was significantly lower in Teff cells (**Figs. 6b, S5a**), in line with our observations from *in vivo* U- [^13^C]glutamine infusions **(Fig. 5b)**. Also, in agreement with U-[^13^C]glutamine infusions, we observed a decrease in total abundance and fractional enrichment of ^13^C-glutamine-derived TCA cycle intermediates (i.e., citrate, α-ketoglutarate, malate) and TCA cycle-derived aspartate in late Teff cells compared to early Teff cells (**Figs. 6c, S5b**).

One prediction is that T cells may switch to glucose to fuel TCA cycle metabolism as glutamine utilization drops late in the T cell response to infection. Through *ex vivo* tracing we found that U-[^13^C]glucose uptake was identical between early and late Teff cells (**Figs. 6d, S5c**). However, processing of glucose through glycolysis was reduced in late Teff cells, with decreased production of 3-phosphoglycerate (3-PG) and lactate from U-[^13^C]-glucose in Teff cells at 6 dpi compared to 3 dpi (**Fig. 6e, S5d**). Notably, early Teff cells displayed glucose-dependent lactate production consistent with the Warburg effect, which was absent in late Teff cells (**Fig. 6e**). Similarly, U-[^13^C]glucose oxidation, like glutaminolysis, was also decreased in late Teff cells compared to early Teff cells. We observed a significant decrease in both the relative abundance (**Fig. 6f**) and fractional enrichment (**Fig. S5e**) of U-[^13^C]glucose carbon in TCA cycle intermediates (i.e., citrate, malate) and aspartate in T cells isolated at late stages of *Lm*OVA infection. Thus, glucose oxidation does not compensate for lower glutamine oxidation at late stages of the T cell response to infection.

Finally, we assessed changes in mitochondrial mass and activity in early and late Teff cell populations. Consistent with lower rates of OXPHOS (**Fig. 5**), late Teff cells displayed lower mitochondrial mass (Mitospy, **Fig. 6g**) and mitochondrial membrane potential (TMRM, **Fig. 6h**) when compared to early Teff cells. This corresponded to a significant decrease in mitochondrial activity per mitochondria (when taking the ratio of TMRM to MitoSpy) (**Fig. 6i**). Taken together, the reduction in OCR (**Fig. 5d**), ATP production capacity (**Fig. 5e**), ^13^C-glutamine- and ^13^C- glucose-derived TCA intermediates (**Figs. 6c, f**), and lower mitochondrial activity (**Fig. 6i**) in late Teff cells indicate that T cells at the peak of the effector response to infection have a reduced bioenergetic profile compared to early effector cells undergoing clonal expansion. Our data suggest that reliance on conventional fuels−specifically glutamine and glucose−correlates with T cell proliferative state (**Fig. 6a**).

### Acetate is physiologic TCA cycle fuel for CD8^+^ T cells

While the contribution of both glucose and glutamine to the TCA cycle in Teff cells declines late in infection (**Fig. 6**), analysis of mass isotopologues of citrate and malate indicate more glutamine incorporation into the TCA cycle than glucose (**Figs. S5b,e**). Specifically, the lack of M+2 labeled TCA intermediates from U-[^13^C]glucose in late Teff cells (**Fig. S5e**) suggests a secondary, non-glucose source of mitochondrial acetyl-CoA contributing to the TCA cycle. We recently reported that several physiologic carbon sources absent from cell culture medium−including β - hydroxybutyrate, lactate, and acetate−can be oxidized by *in vitro*-stimulated CD8^+^ T cells (*18*). As acetate is a metabolic source of acetyl-CoA, we investigated the use of acetate as a TCA cycle fuel *in vivo* via U-[^13^C]acetate infusions. As shown in **Fig. 7a**, we observed low incorporation of U-[^13^C]acetate carbon in TCA cycle intermediates of CD8^+^ T cells at 3 dpi, the stage of infection when U-[^13^C]glutamine contribution to the TCA cycle is highest. However, fractional enrichment of U-[^13^C]acetate-derived carbon in TCA cycle intermediates was increased at the peak of infection (6 dpi) compared to U-[^13^C]glucose and U-[^13^C]glutamine (**Fig. 7a**). This increased labeling at 6 dpi was attributed to both M+2 and M+4 labeling of citrate and malate from U-[^13^C]acetate, indicating that acetate carbon contributed to multiple turns of the TCA cycle *in vivo* (**Fig. 7b**). Additionally, Teff cells at 6 dpi displayed increased fractional enrichment of U-[^13^C]acetate carbon in amino acids generated from TCA cycle metabolism (i.e., glutamate, aspartate, **Fig. 7c**).

**Fig. 7.**
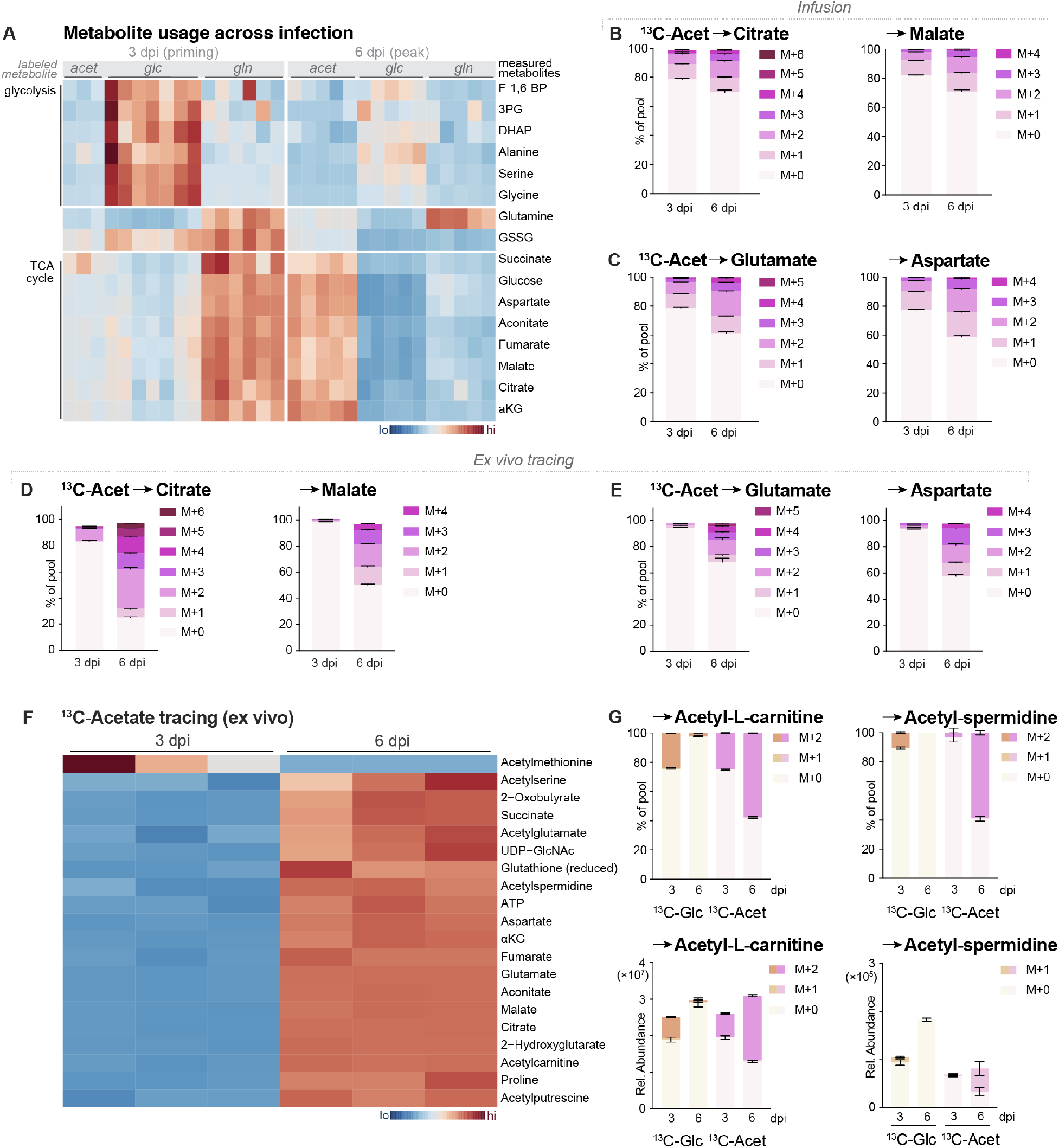
Acetate is physiologic TCA cycle fuel for CD8^+^ T cells. **(A)** Heatmap of relative ^13^C-labeling patterns in intracellular metabolites from Teff cells at early (3 dpi) and peak (6 dpi) stages of *Lm*-OVA infection following 2h infusion of ^13^C-acetate (acet), ^13^C-glucose (glc), or ^13^C-glutamine (gln). Glycolytic and TCA intermediates are highlighted. **(B- C)** ^13^C-acetate tracing into Teff cell metabolites following *in vivo* infusion at 3 or 6 dpi. MID of ^13^C-acetate labeling into intracellular **(B)** TCA cycle intermediates (citrate, malate) and **(C)** TCA cycle-derived amino acids (glutamate, aspartate) in CD8^+^ Teff cells isolated from *Lm*-OVA- infected mice. Mice were infused with ^13^C-acetate for 2h prior to cell isolation. Data represent the mean ± SEM for biological replicates (n=3-6). **(D-E)** ^13^C-acetate incorporation into intracellular metabolites following *ex vivo* culture of CD8^+^ Teff cells with ^13^C-acetate. OT-I CD8^+^ T cells were isolated from *Lm*-OVA-infected mice at 3 or 6 dpi and cultured *ex vivo* for 2h in medium containing 1 mM ^13^C-acetate. Shown are the MID of ^13^C-acetate labeling into intracellular **(D)** citrate and malate or **(E)** glutamate and aspartate in CD8^+^ Teff cells isolated from mice at 3 or 6 dpi. Data represent the mean ± SEM for biological replicates (n=4-5). **(F)** Heatmap of intracellular metabolites in CD8^+^ Teff cells isolated from *Lm*-OVA-infected mice (3 or 6 dpi) displaying enrichment of ^13^C carbon following culture with ^13^C-acetate for 2h *ex vivo*. Shown are metabolites with significant enrichment of ^13^C carbon from ^13^C-acetate (*p*<0.05). **(G)** MID (% of pool) and total abundance of ^13^C-glucose (Glc) or ^13^C-acetate (Acet) labeling into acetylated (M+2) carnitine and spermidine in CD8^+^ Teff cells isolated from mice at 3 or 6 dpi and cultured as in **(D)**.

We recapitulated findings from *in vivo* infusions using *ex vivo* U-[^13^C]acetate tracing of CD8^+^ Teff cells isolated from *Lm*OVA-infected mice at early and late timepoints (3 and 6 dpi, respectively) (**Fig. 7d-e**). This analysis revealed that while acetate was a minor contributor to the overall pool of TCA cycle metabolites early during infection (i.e., less than 20% of the citrate pool at 3 dpi), late Teff cells derived most of their TCA cycle and TCA cycle-derived metabolites from acetate (**Fig. 7d-e**). For example, over 70% of the citrate pool and 50% of the malate pool was derived from extracellular acetate in Teff cells at 6 dpi (**Fig. 7d**), despite similar levels of glucose uptake (**Fig. 6d**). Together these data indicate that acetate is a physiologic TCA cycle fuel for CD8^+^ T cells *in vivo* independent of glucose availability and uptake (*37*).

The overall abundance of metabolic intermediates is lower in late compared to early Teff cells (**Fig. S6a-b**), likely reflecting the loss of glutamine- and glucose-derived carbon in the TCA cycle at later stages of infection. However, focusing on the abundance of U-[^13^C]acetate-derived metabolites in Teff cells over time, the overall amount of acetate-derived TCA cycle metabolites (i.e., citrate and malate, **Fig. S6a**) and amino acids (i.e., glutamate, aspartate, **Fig. S6b**) remains fairly constant. These data argue that acetate contributes carbon to the TCA cycle of Teff cells at all stages of activation but contributes a far smaller fraction to overall TCA cycle metabolism when glucose and glutamine are readily used as fuels. Of note, U-[^13^C]acetate becomes the main TCA cycle fuel source in late Teff cells, as reflected by higher-order labeling patterns (i.e., M+3-6) indicating U-[^13^C]acetate contribution to multiple turns of the TCA cycle in late Teff cells (**Fig. S6c**).

One of the critical functions of mitochondrial-derived citrate is the maintenance of cytosolic acetyl-CoA pools for cellular biosynthesis and acetylation reactions (*38*). Acetate being a prominent source of acetyl-CoA production (*39–41*), we observed enrichment of U-[^13^C]acetate labeling in several acetylated metabolites beyond the TCA cycle in Teff cells (**Fig. 7f**). As a surrogate measurement of acetyl-CoA levels, we assessed the contribution of U-[^13^C]glucose and U-[^13^C]acetate to the acetyl moiety (M+2) of acetylated metabolites in Teff cells over the course of infection. Glucose and acetate contributed equally to acetylation of carnitine and the polyamines spermidine and putrescene−metabolites associated with effector T cell proliferation (*42–44*)−in clonally expanding Teff cells at 3 dpi (**Figs. 7g, S6d**), while other acetylated metabolites (i.e., acetyl-alanine, acetyl-choline) were minimally labeled from either carbon source (**Fig. S6e-f**). Importantly, we observed a shift in acetyl-carnitine and acetyl-spermidine M+2 labeling patterns from U-[^13^C]glucose to U-[^13^C]acetate in late Teff cells (**Fig. 7g**), suggesting that synthesis of these metabolites is maintained from acetate at later stages of infection. These results indicate that acetate is a major source for acetyl-CoA production in Teff cells, particularly at late stages of infection.

## Discussion

Here we have used *in vivo* infusion of ^13^C-labeled metabolites to define the metabolic preferences of effector CD8^+^ T cells as they are responding to infection by a model pathogen (*Lm*) in vivo. This approach allowed us to track the metabolic and bioenergetic tendencies of T cells in the physiologic environment of an immune response. In contrast to our original observations that glucose is primarily used by Teff cells for biosynthetic processes (i.e., nucleotide and amino acid synthesis) *in vivo* (*10*), the data presented here establish that glutamine is both a bioenergetic and biosynthetic substrate for Teff cells. CD8^+^ Teff cells responding to pathogen infection shunt glutamine into the TCA cycle, where it is used to fuel OXPHOS (and ultimately ATP production) and directed towards anabolic processes such as aspartate synthesis. We demonstrate that glutamine-dependent aspartate synthesis is an essential metabolic function of effector T cells, as silencing of *Got1* reduces CD8^+^ T cell expansion and effector function (i.e., IFN-γ production) *in vivo* **(Fig. 4)**. However, kinetic analysis of fuel utilization by Teff cells shows that both glucose and glutamine utilization wanes at the peak of expansion (6 dpi), despite the continuing need for ATP to fuel effector function. In contrast, the short-chain fatty acid (SCFA) acetate becomes a dominant TCA cycle fuel for effector T cells at the peak of the immune response **(Fig. 7).** Together our data highlight a critical role for glutaminolysis fueling early CD8^+^ T cell expansion *in vivo*, with increased plasticity in TCA cycle fuel choice by effector T cells at different stages of infection.

Glutamine is the most abundant circulating amino acid and consumed by cells up to 100- fold more than other amino acids (*45*). Fundamental observations in cancer metabolism have established that glutamine fuels oxidative metabolism for ATP production in a majority of tumor cells (*46, 47*). Similar dependences on glutamine for T cell proliferation and growth *in vitro* have been reported. T cells lacking glutaminase (*Gls*)−the enzyme that generates glutamate from glutamine−expand poorly and display impaired effector function *in vivo* (*48*), while glutaminase inhibitors can reduce the effectiveness of CD8^+^ T cells in the context of anti-tumor immunity (*49*). However, glutamine-derived glutamate production has several potential metabolic fates beyond oxidation, including glutathione production (*50*). Reductive carboxylation of glutamine-derived α-ketoglutarate is prominent alternative metabolic fate for glutamine in cells with impaired mitochondrial oxidative capacity (*33*). Moreover, recent reports establishing that some lung tumor cells with high glutamine requirements *in vitro* do not display glutamine dependency *in vivo* (*26*) have called into question whether glutamine is a bona fide TCA cycle fuel *in vivo*. Our *in vivo* and *ex vivo* tracing data establish that glutamine anaplerosis is a dominant metabolic process in expanding early effector T cells (EECs) *in vivo*, and that early proliferating T cells *in vivo* depend on glutamine to drive OXPHOS.

Circulating glutamine concentrations in mouse plasma are considerably lower than glucose (0.4-0.9 mM, compared to 4-6 mM) (*18*); yet for Teff cells our infusion data indicate that glutamine contributes more carbon to TCA cycle metabolism than glucose *in vivo* (**Fig. 1b**). In addition, reductive carboxylation of α-ketoglutarate is not a metabolic fate of glutamine in Teff cells *in vivo*, even late in infection when mitochondrial membrane potential is lower (**Figs. S5b** and **6h**). We hypothesize that the preferential use of glutamine over glucose to fuel ATP production in EECs is not necessarily governed by metabolite abundance, but rather by optimization of fuel choice matched to cellular function. At 3 dpi, effector T cells are rapidly proliferating and display high demand for both ATP and biosynthetic substrates for growth (i.e., nucleotides, lipids). Our *in vivo* tracing data demonstrate that during this anabolic growth phase glucose is used to produce ribose and amino acids needed for nucleotide synthesis (*10, 11*). In particular, using citrate for lipid synthesis and aspartate for nucleotide synthesis runs the risk of depleting TCA cycle intermediates if additional carbon is not imported to replenish the cycle (*51, 52*). *In vivo*, increased ^13^C-glutamine uptake and input into the TCA cycle solves this issue for Teff cells.

Our *in vivo* tracing data reveal that the metabolic requirements of Teff cells for glutamine extend beyond bioenergetics. We find infused ^13^C-glutamine carbon to be highly enriched in intracellular proline from Teff cells, indicating that in vivo proliferating T cells actively use ^13^C- glutamine-derived glutamate for *de novo* proline synthesis. Similarly, M+3 labeling from ^13^C- glutamine into pyrimidine nucleotides and their derivatives (i.e., UMP, UDP-Glucose) indicates that *de novo* aspartate production is an active anabolic node in Teff cells *in vivo*. Mechanistically, we show that Got1 is a regulator of glutamine-dependent aspartate production in T cells and is required for CD8^+^ T cell responses in vivo. We observed significant decreases in both the expansion and function of antigen-specific CD8^+^ Teff cells in response to *Lm*-OVA infection when *Got1* gene expression was silenced **(Fig. 4F-G).** Recently published work by Xu et al. established similar requirements for Got1 in mediating T cell-mediated anti-tumor responses (*36*). Genetic ablation of *Got1* using T cell-specific conditional deletion in mice (*Got1^fl/fl^ CD4Cre*) resulted in poor control of experimental tumors (i.e., B16 melanoma), due in part to reduced perforin expression and CTL killing capacity by Got1-deficient Teff cells (*36*). It is unclear whether Got2, which is localized in the mitochondria, can functionally compensate for Got1 to maintain intracellular aspartate at sufficient levels to support T cell function. We show that T cells can function normally in the absence of exogenous aspartate; however, *Got1*-silenced T cells become dependent on exogenous aspartate for cell proliferation (**Fig. 4E**). These data suggest that Got1 and Got2 likely work together to buffer T cells against fluctuations in environmental aspartate levels that can impact T cell proliferation.

Current models of T cell metabolism posit a binary switch between catabolic and anabolic metabolism as T cells shift from quiescent (i.e., naïve, memory) to activated (i.e., effector) states, respectively (*53, 54*). In this model, glucose and glutamine are the dominant fuels for effector T cells. The data presented here suggest needed refinements to this model. First, our *in vivo* infusion data clearly support observations from *in vitro* culture models that glucose and glutamine are highly consumed by proliferating T cells. However, these data also indicate that glucose and glutamine label distinct metabolite pools in Teff cells *in vivo*. Glucose is predominantly used for the synthesis of ribose-containing macromolecules (i.e., nucleotides, NAD^+^, SAM/SAH) and glycolysis-derived amino acids (i.e., alanine, serine, glycine), while glutamine provides the bulk of carbon used to power the TCA cycle and produce TCA cycle-derived amino acids (i.e., aspartate) (**Fig. 1B**). Thus, in proliferating T cells glutamine serves as a primary fuel source, leaving glucose to function more as a biosynthetic precursor (*10*).

Second, the use of glutamine and glucose to fuel anabolic metabolism is highly correlated with rates of T cell proliferation. T cell bioenergetic capacity, TCA cycle metabolism, and anabolic biosynthesis are highest in EECs isolated early post infection (2-3 dpi) when CD8^+^ T cells are undergoing the highest rates of proliferation. At the peak of the T cell response to infection (6-7 dpi), Teff cells continue to take up glucose and glutamine at similar rates to EECs but do not use these substrates to fuel TCA cycle metabolism or synthesize biomolecules as they did earlier in the T cell response. This may be due in part to changes in cellular phenotype but most likely due to the shift in proliferative rate and reduced requirements for cellular growth (**Fig. 6A**).

Third, Teff cells at late stages of infection change their preference of oxidative fuel choice, moving away from glutamine and instead using acetate to fuel TCA cycle metabolism (**Fig. 7A**). We recently reported that T cells are capable of oxidizing many different carbon sources−including lactate, β-hydroxybutyrate (βOHB), and acetate−when cultured under physiologic growth conditions (*18*), indicating that activated T cells have a high degree of flexibility when it comes to oxidative fuel choice. While acetate has previously been shown to be consumed by glucose-starved and memory-like T cells *in vitro* (*37, 55, 56*), ours is the first demonstration that Teff cells use acetate as an oxidative fuel *in vivo*. The mechanisms underlying this switch in fuel choice remain unclear. Interestingly, acetate appears to be oxidized by Teff cells at a consistent rate both early and late during infection but proportionally contributes more to the TCA cycle later during infection due to decreased glutamine oxidation at these later timepoints (**Fig. S6A**). Acetate levels have been reported to peak at sites of inflammation and is also produced by the microbiome (*55–57*), which may dictate the preferential use of acetate by T cells simply by being readily available. Alternatively, changes in glutamine availability may dictate the switch to acetate oxidation. Blocking glutamine utilization using the glutamine antagonist 6-diazo-5-oxo-L-norleucine (DON) stimulates increased expression of the acetate-processing enzymes acyl-coenzyme A (CoA) synthetase short-chain family member 1 (ACSS1) and ACSS2, leading to increased acetate utilization by CD8^+^ T cells (*58*). Future work will focus on the impact of acetate metabolism of effector T cell function and cell fate decisions during the late stages of infection.

Together, our findings show T cells display distinct patterns of fuel choice *in vivo* that change over the course of an immune response to infection. These data highlight the importance of investigating the dynamics of effector CD8^+^ T cell metabolic patterns *in vivo*, which may help identify key metabolic nodes that influence T cell effector function and/or differentiation.

## Materials and Methods

### Mice

C57BL/6J, CD90.1 (Thy1.1^+^), and Tg(TcraTcrb)1100Mjb (OT-I) mice were purchased from The Jackson Laboratory (Bar Harbor, ME). Mice were bred and maintained under specific pathogen-free conditions at McGill University and Van Andel Institute under approved protocols. Experiments were performed using mice between 6 and 20 weeks of age. Mice were kept in groups of 5 mice or less and had access to a pellet based feed and autoclaved reverse osmosis water.

### Cell lines

Female 293T (CRL-3216) cells were cultured in Dulbecco’s Modified Eagle’s Medium (DMEM) (Wisent Inc., St. Bruno, QC, Canada) supplemented with 10% heat-inactivated fetal bovine serum (FBS), 1% penicillin-streptomycin (Gibco), and L-glutamine at a final concentration of 6 mM. Cells were cultured at 37°C in a humidified 5% CO_2_ incubator.

### T cell purification and culture

T cells were purified from the spleen and peripheral lymph nodes of C57BL/6 or OT-I mice by negative selection (StemCell Technologies, Vancouver, BC, Canada). Cells were cultured in T cell medium (TCM) containing IMDM or custom IMDM for tracing studies (without glucose, and glutamine) supplemented with 10% dialyzed FBS (Wisent, St. Bruno, QC), L-glutamine (Invitrogen, Chicago, IL), penicillin-streptomycin (Invitrogen), 2-ME (Sigma-Aldrich, St. Louis, MO), and glucose. For *in vitro* activation of OT-I T cells, splenocytes from OT-I mice (1 x 10^6^ cells/ml) were stimulated with OVA_257_ peptide (1 μM) for 2 days, and subsequently expanded in IL2 (50 U/ml, PeproTech, Rocky Hill, NJ) for an additional 2 days. For naïve CD8^+^ mouse T cell cultures, naïve CD8^+^ T cells were purified from spleen and peripheral lymph nodes by negative selection (StemCell Technologies, Vancouver, BC, Canada). Isolated naïve CD8^+^ cells (1 x 10^6^ cells/mL) were cultured in TCM supplemented with 100ng/mL IL-7. CD8^+^ Tn or Teff cells isolated from LmOVA-infected mice by Thy1.1 positive selection kit (StemCell Technologies, Vancouver, BC, Canada) and were cultured immediately *ex vivo* in Seahorse XF medium (non-buffered DMEM 1640 containing 25mM glucose, 2mM L-glutamine, and 1mM sodium pyruvate) or TCM containing 10% dialyzed FBS. For T cell retrovirus transduction CD8^+^ CD90.1^+^ OT-1^+^ T cells were stimulated with plate-bound anti-CD3ε (clone 2C11) and anti-CD28 (clone 37.51) (eBioscience, San Diego, CA) for 24 hours prior to transduction with retrovirus and expanded up 1 day post-transduction in TCM for 2 days in IL2 (50 U/ml). Transduced T cells were sorted by FACS Aria/ BD FACSSymphonyS6 or MoFlo Astrios and sorted T cells were kept in culture overnight before T cell proliferation was measured by cell counts or transduced OT-I T cells transferred into mice.

### Infection with L. monocytogenes

Mice were immunized IV with a sublethal dose of recombinant attenuated *Listeria monocytogenes* expressing OVA (*Lm*OVA, 2 x 10^6^ CFU) as previously described (*27, 59*). For OT-I adaptive transfer experiments (transduced cells), 5 x 10^3^ CD8^+^ OT-I T cells (CD90.1^+^) were injected intravenously into C57BL/6 mice, followed by *Lm*OVA infection 1 day later. Splenocytes were isolated from mice at 7 dpi and analyzed for the presence of OVA-specific CD8^+^ T cells by CD90.1 (for adoptive transfer experiments of transduced T cells). Cytokine production by CD8^+^ T cells was analyzed by ICS staining following peptide re-stimulation (OVA_257_) as previously described (*60*). To generate *in vivo* activated CD8^+^ T cells, for metabolomics, proteomics, and Seahorse bioanalyzer, CD8^+^ Thy1.1^+^ OT-I T cells were injected intravenously into C57BL/6 mice, followed by *Lm*-OVA infection 1 day later. 2 x 10^6^ or 5 x 10^4^ cells were injected and isolated at 3 or 6 dpi respectively.

### Retrovirus production and transduction

For retrovirus production, 293T cells were co-transfected with pCL-Eco and either pLMPd-Amt sh*FF* (control vector) or sh*Got1* using Lipofectamine 2000 transfection reagent (Invitrogen) according to the manufacturer’s protocol. Viral supernatants were harvested 48 and 72 h post-transfection, pooled, and concentrated using Lenti-X Concentrator (Takara Bio) according to the manufacturer’s protocol. The concentrated retrovirus was added to 24 h-activated T cells together with 8 ng/mL polybrene, 200 U/mL IL-2, and 20 mM HEPES, and the cells were centrifuged at 1180 RCF, 30°C for 90 min. Ametrine-positive cells were sorted by FACS 48-72 h post-transduction.

### Flow cytometry, viability, and intracellular cytokine staining

Single-cell suspensions were surface stained with fluorescently conjugated Abs against murine CD4, CD8, CD44, Thy1.1, CD62L, KLRG1, Ki67, and CD127 (eBioscience). Mitochondrial staining was assessed using MitoSpy Green, and TMRM following manufacturer’s protocol. Cell viability was assessed using the Fixable Dye eFluor® 780 (eBioscience) following the manufacturer’s protocol. Intracellular cytokine staining (ICS) for IFN-γ was performed as previously described (Ma et al 2019). Briefly, *Lm*-OVA infected splenocytes at 7 dpi from mice adoptively transferred with retrovirus transduced CD8^+^ Thy1.1^+^ OT-1^+^ T cells were stimulated with OVA_257_ peptide (1ug/mL) *ex vivo* for 3 hr, with Golgi Stop (Fisher Sci) added for the final 3 hr of stimulation, followed by ICS and flow cytometry. Flow cytometry was performed on Cytek Aurora cytometer. Analysis was performed using with FlowJo software (Tree Star).

### Immunoblotting

For immunoblotting, cells were lysed in modified AMPK lysis buffer as previously described (*61*). Cleared lysates were resolved by SDS/PAGE, transferred to nitrocellulose, and proteins detected using primary antibodies to Got1 (NOVUS Biologicals).

### Metabolic assays

T cell oxygen consumption rate (OCR) and extracellular acidification rate (ECAR) were measured using a Seahorse XF96 Extracellular Flux Analyzer following established protocols (*11, 28*). In brief, activated T cells (2 x 10^5^) were plated in poly-D-lysine-coated XF96 plates via centrifugation in XF medium. Cellular bioenergetics were assessed through the sequential addition of oligomycin (2.0 μM), fluoro-carbonyl cyanide phenylhydrazone (FCCP, 2.0 μM), rotenone/antimycin A (2 μM), and monensin (10mM). Data were normalized to cell number. Bioenergetics data analysis was based on protocols developed by Mookerjee and Brand (*62*) and is available for download at https://russelljoneslab.vai.org/tools.

### Stable isotope labeling (SIL) and in vivo ^13^C infusions

SIL experiments with *in vitro*-activated T cells using liquid chromatography (LC) or gas chromatography (GC) coupled to mass spectrometry (MS) were conducted as previously described (*10, 18*). In brief, *in vitro*-activated CD8^+^ T cells or OT-I T cells isolated from *Lm*-OVA-infected mice were washed in IMDM or VIM medium (*18*) containing 10% dialyzed FBS, and re-cultured (2.5 × 10^6^ cells/well in 24-well plates) for 2h in medium containing ^13^C-labeled metabolites at the following concentrations: [U-^13^C_6_]glucose, 5 mM; [U-^13^C_2_]acetate, 1 mM; or [U-^13^C_5_]glutamine,0.5 mM. Cells were transferred from tissue culture plates to falcon tubes and centrifuged at 500 RCF, 4°C for 3 min. The cell pellet was washed with ice-cold saline before being snap frozen on dry ice and stored at -80°C. Metabolites were extracted as described previously (*18*).

*In vivo* infusions in *Lm*-OVA-infected mice were conducted as previously described (*10, 29*). In brief, infected mice were anesthetized using continuous isoflurane exposure and infused for 2h via the tail vein. The following metabolites and delivery rates were used: ^13^C-glucose (100 mg/ml concentration, 120 μL bolus dose, 2.5 μL/min continuous infusion); ^13^C-glutamine (40 mg/ml concentration, 150 μL bolus dose, 3.0 μL/min continuous infusion); ^13^C-acetate (40 mg/ml concentration, 150 μL bolus dose, 3.0 μL/min continuous infusion). At the end of the infusion, blood was collected by cardiac puncture. Following blood collection, a lobe of liver and a piece of spleen were collected and flash frozen in liquid nitrogen and kept at -80°C. The rest of the spleen was turned into a single cell suspension, filtered through a 0.75 μm filter, and pelleted. The cell pellet was resuspended in 4 mL of EasySep buffer (PBS with 2% FBS and 1mM EDTA) and volume split equally for Thy1.1^+^ isolation (EasySep Mouse CD90.1 Positive Selection Kit) and naïve CD8^+^ T cell isolation (EasySep Mouse Naïve CD8+ T cell isolation kit). Cells were isolated as described (*29*) and flash-frozen for metabolomic analysis.

### GC-MS analysis of ^13^C metabolites

Gas chromatography coupled to mass spectrometry (GC-MS) was performed on T cells using previously described methods (*28, 63, 64*). Briefly, activated T cells were washed with PBS and re-cultured in TCM (lacking glucose and serine) containing 10% dialyzed FBS and uniformly labeled [^13^C]-Glucose (Cambridge Isotope Laboratories). T cells (5 x 10^6^ per well in 6 well plates) were cultured in ^13^C-containing medium for up to 6 hours. For cellular media samples, 20 μL of media were taken at indicated time points and centrifuged to remove cells, with 10 μL of media used for metabolite analysis. Metabolites were extracted using ice cold 80% methanol, sonicated, and then D-myristic acid was added (750ng/sample) as an internal standard. Dried samples were dissolved in 30 μL methozyamine hydrochloride (10mg/ml) in pyridine and derivatized as tert-butyldimethylsily (TBDMS) esters using 70 μL N-(*tert*-butyldimethylsilyl)-N-methyltrifluoroacetamide (MTBSTFA) (*61*).

For metabolite analysis, an Agilent 5975C GC/MS equipped with a DB-5MS+DG (30 m x 250 µm x 0.25 µm) capillary column (Agilent J&W, Santa Clara, CA, USA) was used. All data were collected by electron impact set at 70 eV. A total of 1 μL of the derivatized sample was injected in the GC in splitless mode with inlet temperature set to 280°C, using helium as a carrier gas with a flow rate of 1.5512 mL/min (rate at which myristic acid elutes at 17.94 min). The quadrupole was set at 150°C and the GC/MS interface at 285°C. The oven program for all metabolite analyses started at 60°C held for 1 min, then increased at a rate of 10°C/min until 320°C. Bake-out was at 320°C for 10 min. Sample data were acquired both in scan (1-600 m/z) and selected ion monitoring (SIM) modes. Mass isotopomer distribution for cellular metabolites was determined using a custom algorithm developed at McGill University (*65*). Briefly, the atomic composition of the TBDMS-derivatized metabolite fragments (M-57) was determined, and matrices correcting for natural contribution of isotopomer enrichment were generated for each metabolite. After correction for natural abundance, a comparison was made between non-labeled metabolite abundances (^12^C) and metabolite abundances which were synthesized from the ^13^C tracer. Metabolite abundance was expressed relative to the internal standard (D-myristic acid) and normalized to cell number.

### LC-MS analysis of ^13^C metabolites

Metabolites were analyzed for relative abundance by high resolution accurate mass detection (HRAM) on two QExactive™ Orbitrap mass spectrometers (Thermo Fisher Scientific) coupled to Thermo Vanquish liquid chromatography systems as previously described (*18*). Separate instruments were used for negative and positive mode analysis. For negative mode analysis, an Acquity T3 HSS (1.8 µm, 2.1 mm x 150 mm) column (Waters, Eschborn, Germany) was used for chromatographic separation and the elution gradient was carried out with a binary solvent system. Solvent A consisted of 3% methanol, 10 mM tributylamine, and 15mM acetic acid in water (pH 5.0 +/- 0.05) and solvent B was 100% methanol. A constant flow rate of 200 μL min^−1^ was maintained and the linear gradient employed was as follows: 0–2.5 min 100% A, 2.5–5 min increase from 0 to 20% B, 5–7.5 min maintain 80% A and 20% B, 7.5–13 min increase from 20 to 55% B, 13–15.5 min increase from 55 to 95% B, 15.5–18.5 min maintain 5% A and 95% B, 18.5– 19 min decrease from 95 – 0% B, followed by 6 min of re-equilibration at 100% A. The heater temperature was set to 400° C and ion spray voltage was set to 2.75 kV. The column temperature was maintained at 25 °C and sample volumes of 10 µL were injected. A 22-minute full-scan method was used to acquire data with *m/z* scan range from 80 to 1200 and resolution of 70,000. The automatic gain control (AGC) target was set at 1e6 and the maximum injection time was 500 ms. For positive mode analysis, an Atlantis T3 (3 µm, 2.1mm ID × 150mm) column (Waters) was used and the elution gradient was carried out with a binary solvent system Solvent A consisted of 0.1% acetic acid and 0.025% heptafluorobutyric acid in water and solvent B was 100% acetonitrile. A constant flow rate of 400 μL min−1 was maintained and the linear gradient employed was as follows: 0–4 min increase from 0 to 30% B, 4–6 min from 30 to 35% B, 6–6.1 min from 35 to 100% B and hold at 100% B for 5min, followed by 5 min of re-equilibration. The heater temperature was set to 300° C and the ion spray voltage was set to 3.5 kV. The column temperature was maintained at 25 °C and sample volumes of 10 µL were injected. An 11-minute full-scan method was used to acquire data with *m/z* scan range from 70 to 700 and resolution of 70,000. The automatic gain control (AGC) target was set at 1e6 and the maximum injection time was 250 ms. Instrument control and acquisition was carried out by Xcalibur 2.2 software (Thermo Fisher Scientific).

### GC-MS/LC-MS metabolite identification and relative quantification

Full scan GC-MS/LC-MS data was centroided using vendor software, converted to mzXML format, and further analyzed using customized open-source software (El-Maven). Compounds were identified by *m/z* and retention time: expected *m/z* of de-protonated species was computed based on exact monoisotopic mass, and retention times were matched to those of previously analyzed pure standards and concurrently analyzed control samples. Peak intensities were calculated as the average of the three scans around the peak apex (“AreaTop”).

### Statistical analysis

Data are presented as mean ± SD for technical replicates or mean ± SEM for biological replicates and were analyzed using unpaired Student’s t test or One-Way ANOVA. Statistical significance is indicated in all figures by the following annotations: *, *p* < 0.05; **, *p* < 0.001; ***, *p* < 0.0001.

## Acknowledgments

We acknowledge Drs. Ralph DeBerardinis, Brandon Faubert, Julian Lum, Sara Nowinski, Evan Lien, Carolyn Anderson, scientists at Agios Pharmaceuticals, and members of the Jones and Krawczyk laboratories for scientific discussions contributing to this manuscript. We thank Teresa Leone, Jeanie Wedberg, Margene Brewer, and Michelle Minard for administrative assistance. We thank members of the VAI Core Facilities (Mass Spectrometry, Bioinformatics and Biostatistics, Flow Cytometry, and Vivarium) for technical assistance.

## Author contributions

Conceptualization: EHM, TR, RGJ

Methodology: EHM, MJV, TR, RDS, RGJ

Investigation: EHM, MSD, LMD, IK, SMKG, DGR, CAS, MJV, RMJ, BS, MS, RDS

Visualization: KSW, MV, RMJ

Supervision: TR, CMK, KSW, RDS, RGJ

Writing—original draft: EHM, MSD, KSW, RGJ

Writing—review & editing: EHM, MSD, KSW, RGJ

## Data and materials availability

All unique/stable reagents generated in this study will be made available from the Lead Contact with a completed Materials Transfer Agreement. Plasmids generated in this study will be deposited to Addgene. All the data are available in the main text or supplemental figures and tables.

## Supplementary Materials

Supplementary figures S1-S6 are available as a PDF file. Raw metabolomics data from infusion studies will be made available as supplementary tables.

## References

1. S. M. Kaech, W. Cui, Transcriptional control of effector and memory CD8+ T cell differentiation. Nat. Rev. Immunol. 12, 749–761 (2012).

2. J. T. Chang, E. J. Wherry, A. W. Goldrath, Molecular regulation of effector and memory T cell differentiation. Nat. Immunol. 15, 1104–1115 (2014).

3. M. D. Buck, D. O’Sullivan, E. L. Pearce, T cell metabolism drives immunity. J. Cell Biol. 210, 2104OIA169 (2015).

4. E. L. Pearce, E. J. Pearce, Metabolic pathways in immune cell activation and quiescence. Immunity. 38, 633–643 (2013).

5. E. L. Pearce, M. C. Poffenberger, C.-H. Chang, R. G. Jones, Fueling immunity: insights into metabolism and lymphocyte function. Science. 342, 1242454 (2013).

6. I. Kaymak, K. S. Williams, J. R. Cantor, R. G. Jones, Immunometabolic Interplay in the Tumor Microenvironment. Cancer Cell (2020), doi:10.1016/j.ccell.2020.09.004.

7. K. Voss, H. S. Hong, J. E. Bader, A. Sugiura, C. A. Lyssiotis, J. C. Rathmell, A guide to interrogating immunometabolism. Nat. Rev. Immunol. 21, 637–652 (2021).

8. M. G. Vander Heiden, L. C. Cantley, C. B. Thompson, Understanding the Warburg effect: the metabolic requirements of cell proliferation. Science. 324, 1029–1033 (2009).

9. K. E. Wellen, G. Hatzivassiliou, U. M. Sachdeva, T. V. Bui, J. R. Cross, C. B. Thompson, ATP-citrate lyase links cellular metabolism to histone acetylation. Science. 324, 1076–1080 (2009).

10. E. H. Ma, M. J. Verway, R. M. Johnson, D. G. Roy, M. Steadman, S. Hayes, K. S. Williams, R. D. Sheldon, B. Samborska, P. A. Kosinski, H. Kim, T. Griss, B. Faubert, S. A. Condotta, C. M. Krawczyk, R. J. DeBerardinis, K. M. Stewart, M. J. Richer, V. Chubukov, T. P. Roddy, R. G. Jones, Metabolic Profiling Using Stable Isotope Tracing Reveals Distinct Patterns of Glucose Utilization by Physiologically Activated CD8+ T Cells. Immunity (2019), doi:10.1016/j.immuni.2019.09.003.

11. E. H. Ma, G. Bantug, T. Griss, S. Condotta, R. M. Johnson, B. Samborska, N. Mainolfi, V. Suri, H. Guak, M. L. Balmer, M. J. Verway, T. C. Raissi, H. Tsui, G. Boukhaled, S. Henriques da Costa, C. Frezza, C. M. Krawczyk, A. Friedman, M. Manfredi, M. J. Richer, C. Hess, R. G. Jones, Serine is an essential metabolite for effector T cell expansion. Cell Metab. 25, 345– 357 (2017).

12. L. K. Smith, G. M. Boukhaled, S. A. Condotta, S. Mazouz, J. J. Guthmiller, R. Vijay, N. S. Butler, J. Bruneau, N. H. Shoukry, C. M. Krawczyk, M. J. Richer, Interleukin-10 directly inhibits CD8+ T cell function by enhancing N-glycan branching to decrease antigen sensitivity. Immunity. 48, 299–312.e5 (2018).

13. M. S. Pereira, I. Alves, M. Vicente, A. Campar, M. C. Silva, N. A. Padrão, V. Pinto, Â. Fernandes, A. M. Dias, S. S. Pinho, Glycans as key checkpoints of T cell activity and function. Front. Immunol. 9, 2754 (2018).

14. N. Ron-Harel, D. Santos, J. M. Ghergurovich, P. T. Sage, A. Reddy, S. B. Lovitch, N. Dephoure, F. K. Satterstrom, M. Sheffer, J. B. Spinelli, S. Gygi, J. D. Rabinowitz, A. H. Sharpe, M. C. Haigis, Mitochondrial Biogenesis and Proteome Remodeling Promote One-Carbon Metabolism for T Cell Activation. Cell Metab. (2016), doi:10.1016/j.cmet.2016.06.007.

15. A. N. Macintyre, V. A. Gerriets, A. G. Nichols, R. D. Michalek, M. C. Rudolph, D. Deoliveira, S. M. Anderson, E. D. Abel, B. J. Chen, L. P. Hale, J. C. Rathmell, The glucose transporter Glut1 is selectively essential for CD4 T cell activation and effector function. Cell Metab. 20, 61–72 (2014).

16. S. M. Hochrein, H. Wu, M. Eckstein, L. Arrigoni, J. S. Herman, F. Schumacher, C. Gerecke, M. Rosenfeldt, D. Grün, B. Kleuser, G. Gasteiger, W. Kastenmüller, B. Ghesquière, J. Van den Bossche, E. D. Abel, M. Vaeth, The glucose transporter GLUT3 controls T helper 17 cell responses through glycolytic-epigenetic reprogramming. Cell Metab. 34, 516–532.e11 (2022).

17. K. Xu, N. Yin, M. Peng, E. G. Stamatiades, A. Shyu, P. Li, X. Zhang, M. H. Do, Z. Wang, K. J. Capistrano, C. Chou, A. G. Levine, A. Y. Rudensky, M. O. Li, Glycolysis fuels phosphoinositide 3-kinase signaling to bolster T cell immunity. Science. 371, 405–410 (2021).

18. I. Kaymak, K. M. Luda, L. R. Duimstra, E. H. Ma, J. Longo, M. S. Dahabieh, B. Faubert, B. M. Oswald, M. J. Watson, S. M. Kitchen-Goosen, L. M. DeCamp, S. E. Compton, Z. Fu, R. J. DeBerardinis, K. S. Williams, R. D. Sheldon, R. G. Jones, Carbon source availability drives nutrient utilization in CD8+ T cells. Cell Metab. 34, 1298–1311.e6 (2022).

19. L. Long, J. Wei, S. A. Lim, J. L. Raynor, H. Shi, J. P. Connelly, H. Wang, C. Guy, B. Xie, N. M. Chapman, G. Fu, Y. Wang, H. Huang, W. Su, J. Saravia, I. Risch, Y.-D. Wang, Y. Li, M. Niu, Y. Dhungana, A. Kc, P. Zhou, P. Vogel, J. Yu, S. M. Pruett-Miller, J. Peng, H. Chi, CRISPR screens unveil signal hubs for nutrient licensing of T cell immunity. Nature. 600, 308–313 (2021).

20. H. Zeng, K. Yang, C. Cloer, G. Neale, P. Vogel, H. Chi, mTORC1 couples immune signals and metabolic programming to establish T(reg)-cell function. Nature. 499, 485–490 (2013).

21. J. D. Powell, G. M. Delgoffe, The mammalian target of rapamycin: linking T cell differentiation, function, and metabolism. Immunity. 33, 301–311 (2010).

22. J. R. Cantor, M. Abu-Remaileh, N. Kanarek, E. Freinkman, X. Gao, A. Louissaint Jr, C. A. Lewis, D. M. Sabatini, Physiologic medium rewires cellular metabolism and reveals uric acid as an endogenous inhibitor of UMP synthase. Cell. 169, 258–272.e17 (2017).

23. J. Vande Voorde, T. Ackermann, N. Pfetzer, D. Sumpton, G. Mackay, G. Kalna, C. Nixon, K. Blyth, E. Gottlieb, S. Tardito, Improving the metabolic fidelity of cancer models with a physiological cell culture medium. Sci. Adv. 5, eaau7314 (2019).

24. S. Hui, A. J. Cowan, X. Zeng, L. Yang, T. TeSlaa, X. Li, C. Bartman, Z. Zhang, C. Jang, L. Wang, W. Lu, J. Rojas, J. Baur, J. D. Rabinowitz, Quantitative fluxomics of circulating metabolites. Cell Metab. 32, 676–688.e4 (2020).

25. M. Leney-Greene, A. Boddapati, H. Su, J. Cantor, M. Lenardo, Human plasma-like medium improves T lymphocyte activation. iScience (2019), doi:10.1101/740845.

26. S. M. Davidson, T. Papagiannakopoulos, B. A. Olenchock, J. E. Heyman, M. A. Keibler, A. Luengo, M. R. Bauer, A. K. Jha, J. P. O’Brien, K. A. Pierce, D. Y. Gui, L. B. Sullivan, T. M. Wasylenko, L. Subbaraj, C. R. Chin, G. Stephanopolous, B. T. Mott, T. Jacks, C. B. Clish, M. G. Vander Heiden, Environment Impacts the Metabolic Dependencies of Ras-Driven Non-Small Cell Lung Cancer. Cell Metab. 23, 517–528 (2016).

27. C.-H. Chang, J. D. Curtis, L. B. Maggi Jr, B. Faubert, A. V. Villarino, D. O’Sullivan, S. C.-C. Huang, G. J. W. van der Windt, J. Blagih, J. Qiu, J. D. Weber, E. J. Pearce, R. G. Jones, E. L. Pearce, Posttranscriptional control of T cell effector function by aerobic glycolysis. Cell. 153, 1239–1251 (2013).

28. J. Blagih, F. Coulombe, E. E. Vincent, F. Dupuy, G. Galicia-Vazquez, E. Yurchenko, T. C. Raissi, G. J. W. van der Windt, B. Viollet, E. L. Pearce, J. Pelletier, C. A. Piccirillo, C. M. Krawczyk, M. Divangahi, R. G. Jones, G. Galicia-Vázquez, E. Yurchenko, T. C. Raissi, G. J. W. van der Windt, B. Viollet, E. L. Pearce, J. Pelletier, C. A. Piccirillo, C. M. Krawczyk, M. Divangahi, R. G. Jones, The energy sensor AMPK regulates T cell metabolic adaptation and effector responses in vivo. Immunity. 42, 41–54 (2015).

29. R. D. Sheldon, E. H. Ma, L. M. DeCamp, K. S. Williams, R. G. Jones, Interrogating in vivo T-cell metabolism in mice using stable isotope labeling metabolomics and rapid cell sorting. Nat. Protoc. (2021), doi:10.1038/s41596-021-00586-2.

30. A. Muir, L. V. Danai, M. G. Vander Heiden, Microenvironmental regulation of cancer cell metabolism: implications for experimental design and translational studies. Dis. Model. Mech. 11 (2018), doi:10.1242/dmm.035758.

31. M. O. Johnson, P. J. Siska, D. C. Contreras, J. C. Rathmell, Nutrients and the microenvironment to feed a T cell army. Semin. Immunol. 28, 505–513 (2016).

32. A. A. Cluntun, M. J. Lukey, R. A. Cerione, J. W. Locasale, Glutamine Metabolism in Cancer: Understanding the Heterogeneity. Trends Cancer Res. 3, 169–180 (2017).

33. A. R. Mullen, W. W. Wheaton, E. S. Jin, P.-H. Chen, L. B. Sullivan, T. Cheng, Y. Yang, W. M. Linehan, N. S. Chandel, R. J. DeBerardinis, Reductive carboxylation supports growth in tumour cells with defective mitochondria. Nature. 481, 385–388 (2011).

34. C. M. Metallo, P. A. Gameiro, E. L. Bell, K. R. Mattaini, J. Yang, K. Hiller, C. M. Jewell, Z. R. Johnson, D. J. Irvine, L. Guarente, J. K. Kelleher, M. G. Vander Heiden, O. Iliopoulos, G. Stephanopoulos, Reductive glutamine metabolism by IDH1 mediates lipogenesis under hypoxia. Nature. 481, 380–384 (2011).

35. N. Chandel, Navigating Metabolism (Cold Spring Harbor Laboratory Press, New York, NY, 2014).

36. W. Xu, C. H. Patel, L. Zhao, I.-H. Sun, M.-H. Oh, I.-M. Sun, R. S. Helms, J. Wen, J. D. Powell, GOT1 regulates CD8+ effector and memory T cell generation. Cell Rep. 42, 111987 (2023).

37. J. Qiu, M. Villa, D. E. Sanin, M. D. Buck, D. O’Sullivan, R. Ching, M. Matsushita, K. M. Grzes, F. Winkler, C. H. Chang, J. D. Curtis, R. L. Kyle, N. Van Teijlingen Bakker, M. Corrado, F. Haessler, F. Alfei, J. Edwards-Hicks, L. B. Maggi, D. Zehn, T. Egawa, B. Bengsch, R. I. Klein Geltink, T. Jenuwein, E. J. Pearce, E. L. Pearce, Acetate Promotes T Cell Effector Function during Glucose Restriction. Cell Rep. (2019), doi:10.1016/j.celrep.2019.04.022.

38. A. Kinnaird, S. Zhao, K. E. Wellen, E. D. Michelakis, Metabolic control of epigenetics in cancer. Nat. Rev. Cancer. 16, 694–707 (2016).

39. S. Zhao, A. M. Torres, R. A. Henry, S. Trefely, M. Wallace, J. V. Lee, A. Carrer, A. Sengupta, S. L. Campbell, Y. M. Kuo, A. J. Frey, N. Meurs, J. M. Viola, I. A. Blair, A. M. Weljie, C. M. Metallo, N. W. Snyder, A. J. Andrews, K. E. Wellen, ATP-Citrate Lyase Controls a Glucose-to-Acetate Metabolic Switch. Cell Rep. (2016), doi:10.1016/j.celrep.2016.09.069.

40. Z. T. Schug, B. Peck, D. T. Jones, Q. Zhang, S. Grosskurth, I. S. Alam, L. M. Goodwin, E. Smethurst, S. Mason, K. Blyth, L. McGarry, D. James, E. Shanks, G. Kalna, R. E. Saunders, M. Jiang, M. Howell, F. Lassailly, M. Z. Thin, B. Spencer-Dene, G. Stamp, N. J. F. van den Broek, G. Mackay, V. Bulusu, J. J. Kamphorst, S. Tardito, D. Strachan, A. L. Harris, E. O. Aboagye, S. E. Critchlow, M. J. O. Wakelam, A. Schulze, E. Gottlieb, Acetyl-CoA synthetase 2 promotes acetate utilization and maintains cancer cell growth under metabolic stress. Cancer Cell. 27, 57–71 (2015).

41. S. A. Comerford, Z. Huang, X. Du, Y. Wang, L. Cai, A. K. Witkiewicz, H. Walters, M. N. Tantawy, A. Fu, H. C. Manning, J. D. Horton, R. E. Hammer, S. L. McKnight, B. P. Tu, Acetate dependence of tumors. Cell. 159, 1591–1602 (2014).

42. D. J. Puleston, F. Baixauli, D. E. Sanin, J. Edwards-Hicks, M. Villa, A. M. Kabat, M. M. Kamiński, M. Stanckzak, H. J. Weiss, K. M. Grzes, K. Piletic, C. S. Field, M. Corrado, F. Haessler, C. Wang, Y. Musa, L. Schimmelpfennig, L. Flachsmann, G. Mittler, N. Yosef, V. K. Kuchroo, J. M. Buescher, S. Balabanov, E. J. Pearce, D. R. Green, E. L. Pearce, Polyamine metabolism is a central determinant of helper T cell lineage fidelity. Cell. 184, 4186–4202.e20 (2021).

43. R. Wang, C. P. P. Dillon, L. Z. Z. Shi, S. Milasta, R. Carter, D. Finkelstein, L. L. L. McCormick, P. Fitzgerald, H. Chi, J. Munger, D. R. R. Green, The Transcription Factor Myc Controls Metabolic Reprogramming upon T Lymphocyte Activation. Immunity. 35, 871–882 (2011).

44. D. O’Sullivan, G. J. W. van der Windt, S. C.-C. Huang, J. D. Curtis, C.-H. Chang, M. D. Buck, J. Qiu, A. M. Smith, W. Y. Lam, L. M. DiPlato, F.-F. Hsu, M. J. Birnbaum, E. J. Pearce, E. L. Pearce, Memory CD8(+) T cells use cell-intrinsic lipolysis to support the metabolic programming necessary for development. Immunity. 41, 75–88 (2014).

45. H. Eagle, The minimum vitamin requirements of the L and HeLa cells in tissue culture, the production of specific vitamin deficiencies, and their cure. J. Exp. Med. 102, 595–600 (1955).

46. J. Fan, J. J. Kamphorst, R. Mathew, M. K. Chung, E. White, T. Shlomi, J. D. Rabinowitz, Glutamine-driven oxidative phosphorylation is a major ATP source in transformed mammalian cells in both normoxia and hypoxia. Mol. Syst. Biol. 9, 712 (2013).

47. R. J. DeBerardinis, A. Mancuso, E. Daikhin, I. Nissim, M. Yudkoff, S. Wehrli, C. B. Thompson, Beyond aerobic glycolysis: transformed cells can engage in glutamine metabolism that exceeds the requirement for protein and nucleotide synthesis. Proc. Natl. Acad. Sci. U. S. A. 104, 19345–19350 (2007).

48. M. O. Johnson, M. M. Wolf, M. Z. Madden, G. Andrejeva, A. Sugiura, D. C. Contreras, D. Maseda, M. V. Liberti, K. Paz, R. J. Kishton, M. E. Johnson, A. A. de Cubas, P. Wu, G. Li, Y. Zhang, D. C. Newcomb, A. D. Wells, N. P. Restifo, W. K. Rathmell, J. W. Locasale, M. L. Davila, B. R. Blazar, J. C. Rathmell, Distinct Regulation of Th17 and Th1 Cell Differentiation by Glutaminase-Dependent Metabolism. Cell (2018), doi:10.1016/j.cell.2018.10.001.

49. S. A. Best, P. M. Gubser, S. Sethumadhavan, A. Kersbergen, Y. L. Negrón Abril, J. Goldford, K. Sellers, W. Abeysekera, A. L. Garnham, J. A. McDonald, C. E. Weeden, D. Anderson, D. Pirman, T. P. Roddy, D. J. Creek, A. Kallies, G. Kingsbury, K. D. Sutherland, Glutaminase inhibition impairs CD8 T cell activation in STK11-/Lkb1-deficient lung cancer. Cell Metab. 34, 874–887.e6 (2022).

50. W.-K. Chang, K. D. Yang, H. Chuang, J.-T. Jan, M.-F. Shaio, Glutamine protects activated human T cells from apoptosis by up-regulating glutathione and Bcl-2 levels. Clin. Immunol. 104, 151–160 (2002).

51. R. J. DeBerardinis, J. J. Lum, G. Hatzivassiliou, C. B. Thompson, The biology of cancer: metabolic reprogramming fuels cell growth and proliferation. Cell Metab. 7, 11–20 (2008).

52. L. Yang, S. Venneti, D. Nagrath, Glutaminolysis: A Hallmark of Cancer Metabolism. Annu. Rev. Biomed. Eng. 19, 163–194 (2017).

53. N. M. Chapman, H. Chi, Metabolic adaptation of lymphocytes in immunity and disease. Immunity. 55, 14–30 (2022).

54. N. M. Chapman, M. R. Boothby, H. Chi, Metabolic coordination of T cell quiescence and activation. Nat. Rev. Immunol. 20, 55–70 (2020).

55. M. L. Balmer, E. H. Ma, G. R. Bantug, J. Grählert, S. Pfister, T. Glatter, A. Jauch, S. Dimeloe, E. Slack, P. Dehio, M. A. Krzyzaniak, C. G. King, A.-V. Burgener, M. Fischer, L. Develioglu, R. Belle, M. Recher, W. V. Bonilla, A. J. Macpherson, S. Hapfelmeier, R. G. Jones, C. Hess, Memory CD8(+) T cells require increased concentrations of acetate induced by stress for optimal function. Immunity. 44, 1312–1324 (2016).

56. M. L. Balmer, E. H. Ma, A. J. Thompson, R. Epple, G. Unterstab, J. Lötscher, P. Dehio, C. M. Schürch, J. D. Warncke, G. Perrin, A.-K. Woischnig, J. Grählert, J. Löliger, N. Assmann, G. R. Bantug, O. P. Schären, N. Khanna, A. Egli, L. Bubendorf, K. Rentsch, S. Hapfelmeier, R. G. Jones, C. Hess, Memory CD8+ T cells balance pro- and anti-inflammatory activity by reprogramming cellular acetate handling at sites of infection. Cell Metab. 32, 457–467.e5 (2020).

57. M. K. De Siqueira, V. Andrade-Oliveira, A. Stacy, J. Pedro Tôrres Guimarães, R. Wesley Alberca-Custodio, A. Castoldi, J. Marques Santos, M. Davoli-Ferreira, L. Menezes-Silva, W. Miguel Turato, S.-J. Han, A. Glatman Zaretsky, T. W. Hand, N. Olsen Saraiva Câmara, M. Russo, S. Jancar, D. Morais da Fonseca, Y. Belkaid, Infection-elicited microbiota promotes host adaptation to nutrient restriction. Proc. Natl. Acad. Sci. U. S. A. 120, e2214484120 (2023).

58. R. D. Leone, L. Zhao, J. M. Englert, I. M. Sun, M. H. Oh, I. H. Sun, M. L. Arwood, I. A. Bettencourt, C. H. Patel, J. Wen, A. Tam, R. L. Blosser, E. Prchalova, J. Alt, R. Rais, B. S. Slusher, J. D. Powell, Glutamine blockade induces divergent metabolic programs to overcome tumor immune evasion. Science (2019), doi:10.1126/science.aav2588.

59. C. M. Krawczyk, H. Shen, E. J. Pearce, Memory CD4 T cells enhance primary CD8 T-cell responses. Infect. Immun. 75, 3556–3560 (2007).

60. R. G. Jones, T. Bui, C. White, M. Madesh, C. M. Krawczyk, T. Lindsten, B. J. Hawkins, S. Kubek, K. A. Frauwirth, Y. L. Wang, S. J. Conway, H. L. Roderick, M. D. Bootman, H. Shen, J. K. Foskett, C. B. Thompson, The proapoptotic factors Bax and Bak regulate T Cell proliferation through control of endoplasmic reticulum Ca(2+) homeostasis. Immunity. 27, 268–280 (2007).

61. B. Faubert, E. E. Vincent, T. Griss, B. Samborska, S. Izreig, R. U. Svensson, O. A. Mamer, D. Avizonis, D. B. Shackelford, R. J. Shaw, R. G. Jones, Loss of the tumor suppressor LKB1 promotes metabolic reprogramming of cancer cells via HIF-1α. Proc. Natl. Acad. Sci. U. S. A. 111, 2554–2559 (2014).

62. S. A. Mookerjee, D. G. Nicholls, M. D. Brand, Determining maximum glycolytic capacity using extracellular flux measurements. PLoS One. 11, e0152016 (2016).

63. T. Griss, E. E. Vincent, R. Egnatchik, J. Chen, E. H. Ma, B. Faubert, B. Viollet, R. J. DeBerardinis, R. G. Jones, Metformin antagonizes cancer cell proliferation by suppressing mitochondrial-dependent biosynthesis. PLoS Biol. 13, e1002309 (2015).

64. E. E. Vincent, A. Sergushichev, T. Griss, M.-C. Gingras, B. Samborska, T. Ntimbane, P. P. Coelho, J. Blagih, T. C. Raissi, L. Choinière, G. Bridon, E. Loginicheva, B. R. Flynn, E. C. Thomas, J. M. Tavaré, D. Avizonis, A. Pause, D. J. E. Elder, M. N. Artyomov, R. G. Jones, Mitochondrial phosphoenolpyruvate carboxykinase regulates metabolic adaptation and enables glucose-independent tumor growth. Mol. Cell. 60, 195–207 (2015).

65. S. McGuirk, S.-P. Gravel, G. Deblois, D. J. Papadopoli, B. Faubert, A. Wegner, K. Hiller, D. Avizonis, U. D. Akavia, R. G. Jones, V. Giguère, J. St-Pierre, PGC-1α supports glutamine metabolism in breast cancer. Cancer Metab. 1, 22 (2013)yyyyyyyyyyyyyyyyyyy.

