## Supporting Material for "^13^C metabolite tracing reveals glutamine and acetate as critical in vivo fuels for CD8^+^ T cells"

Science Advances
AAAS

### Supplementary Materials for

#### $^{13}C$ metabolite tracing reveals glutamine and acetate as critical in vivo fuels for $CD8^+$ T cells

Eric H. Ma *et al.*

**This PDF file includes:**

Figs. S1 to S6

**Fig. S1**

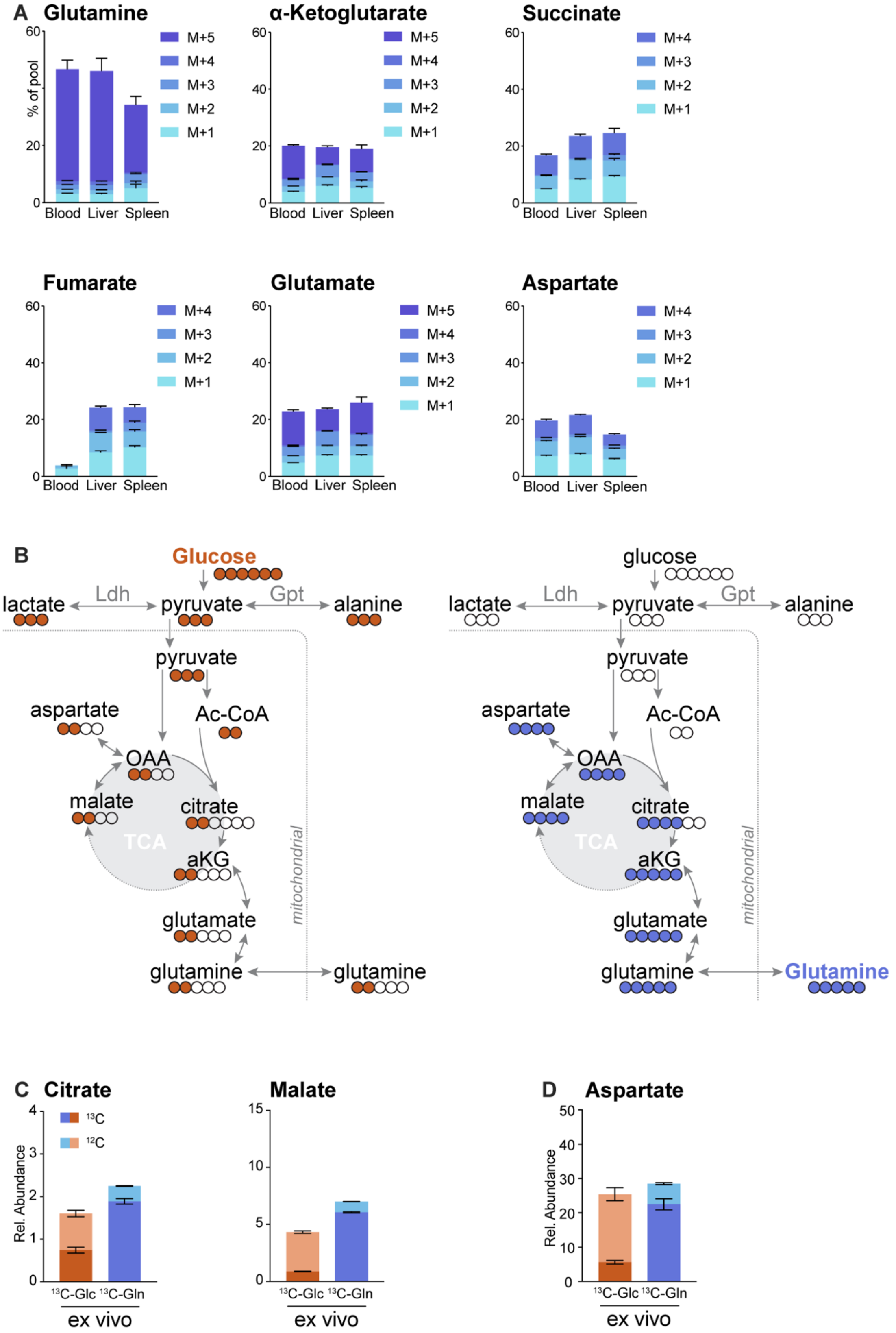
A
Glutamine
α-Ketoglutarate
Succinate
Fumarate
Glutamate
Aspartate
% of pool
M+5
M+4
M+3
M+2
M+1
Blood Liver Spleen
0
20
40
60
B
Glucose
glucose
lactate
Ldh
pyruvate
Gpt
alanine
Ac-CoA
aspartate
OAA
malate
citrate
TCA
aKG
glutamate
glutamine
Glutamine
mitochondrial
C
Citrate
Malate
$^{13}$C
$^{12}$C
Rel. Abundance
$^{13}$C-Glc $^{13}$C-Gln
ex vivo
D
Aspartate

**Fig. S1. U-[$^{13}$C]glucose and U-[$^{13}$C]glutamine labeling patterns in tissues and T cells during *Lm*-OVA infection. (A)** Tissue mass isotopologue distribution (MID) patterns of TCA intermediates upon U-[$^{13}$C]glutamine infusion. MID of $^{13}$C-glutamine labeling (% of pool) in glutamine, TCA cycle intermediates (α-ketoglutarate, succinate, fumarate), and TCA cycle-derived amino acids (glutamate, aspartate) in blood, liver, and spleen in *Lm*-OVA infected mice (3 dpi) following 2h infusion of U-[$^{13}$C]glutamine. Data represent the mean ± SEM for biological replicates (n=6). **(B)** Schematic depicting the metabolic fates of U-[$^{13}$C]glucose (left) and U-[$^{13}$C]glutamine (right). Closed circles depict heavy-labeled ($^{13}$C) carbon atoms in metabolic intermediates. Legend: *TCA,* Tricarboxylic acid cycle (TCA); Ldh*,* Lactate dehydrogenase; Gpt*,* Glutamate-Pyruvate Transaminase. **(C-D)** Relative abundance of $^{12}$C- versus $^{13}$C-labeled intracellular **(C)** citrate and malate, and **(D)** aspartate in CD8$^{+}$ Teff cells isolated from *Lm*-OVA-infected mice (3 dpi). Isolated T cells were cultured for 2h *ex vivo* with $^{13}$C-Glc or $^{13}$C-Gln prior to metabolite extraction. Data represent the mean ± SEM for biological replicates (n=3).

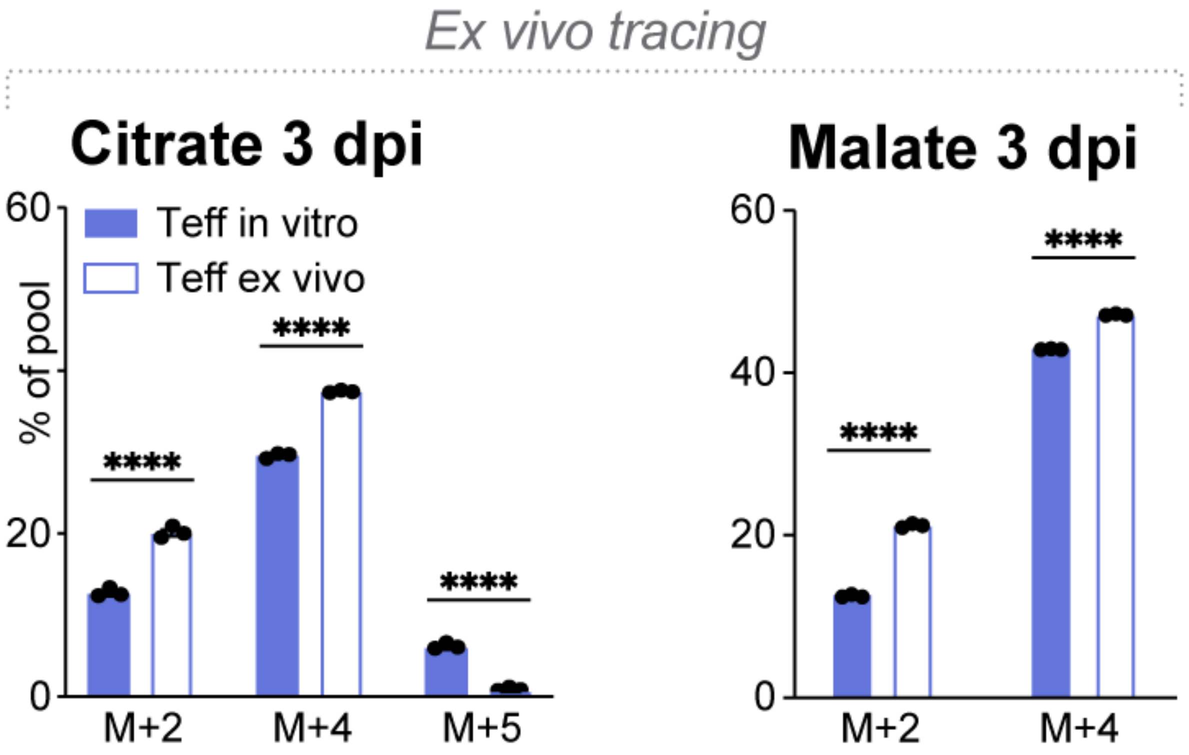


**Fig. S2. Comparison of $^{13}$C-Glutamine labeling patterns in TCA cycle intermediates between *in vitro*- versus *in vivo*-activated T cells.** MID for U-[$^{13}$C]glutamine labeling (% of pool) in intracellular citrate and malate in OT-I T cells activated *in vitro* for 3 days or isolated from *Lm*-OVA-infected mice (3 dpi). Cells were cultured for 2h with U-[$^{13}$C]glutamine prior to metabolite extraction. Data represent the mean ± SEM for biological replicates (n=3).

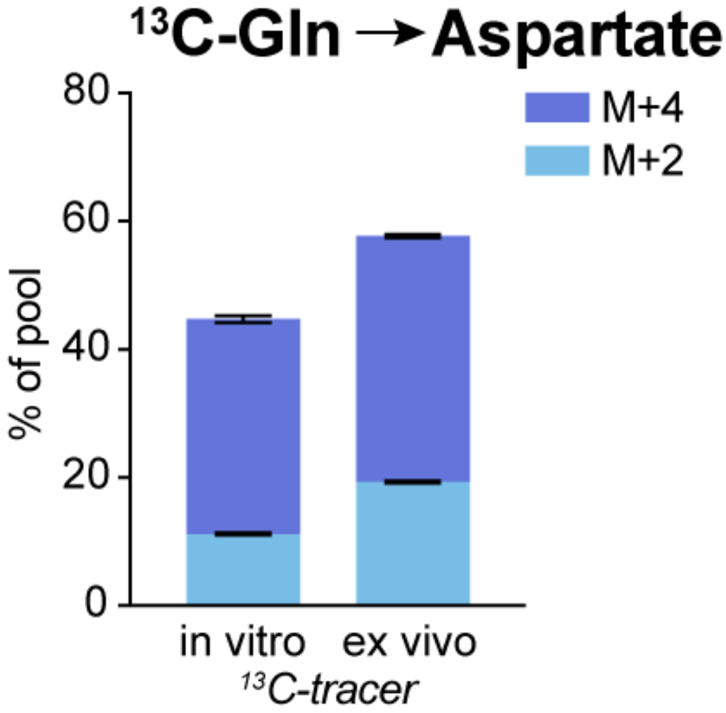


**Fig. S3. U-[$^{13}$C]glutamine labeling into intracellular aspartate in CD8$^+$ T cells.** MID for U-[$^{13}$C]glutamine labeling (% of pool) in intracellular aspartate in OT-I T cells activated *in vitro* for 3 days or isolated from *Lm*-OVA-infected mice (3 dpi). Cells were cultured for 2h with U-[$^{13}$C]glutamine prior to metabolite extraction. Shown are M+2 and M+4 isotopologues of aspartate, plotted as the percentage of total pool. Data represent the mean ± SEM for biological replicates (n=3).

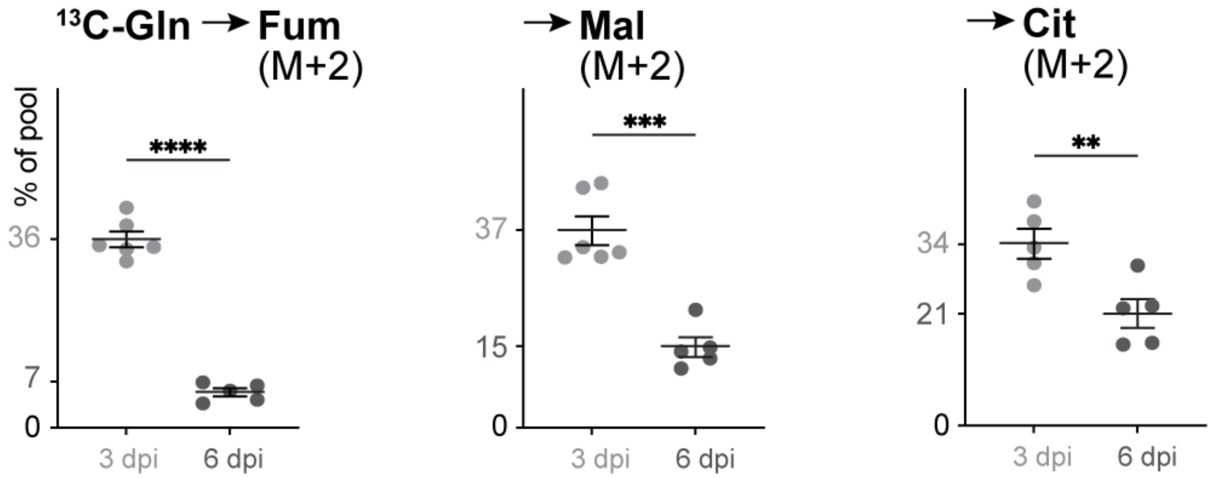


**Fig. S4. Comparison of $^{13}$C-Glutamine labeling patterns in TCA cycle intermediates in T cells following infusion at different stages of infection.** Percent U-[$^{13}$C]glutamine labeling (M+2) in intracellular fumarate (fum), malate (mal) and citrate (cit) in OT-I T cells isolated from *Lm*-OVA-infected mice at 3 and 6 dpi (mean ± SEM, n=5-6). Mice were infused with U-[$^{13}$C]glutamine for 2h prior to metabolite extraction. Data are normalized to serum $^{13}$C-Gln (M+5) levels. M+2 isotopologues designate the second turn of the TCA cycle while M+4 (see Figure 5) indicates the first turn.

Fig. S5.

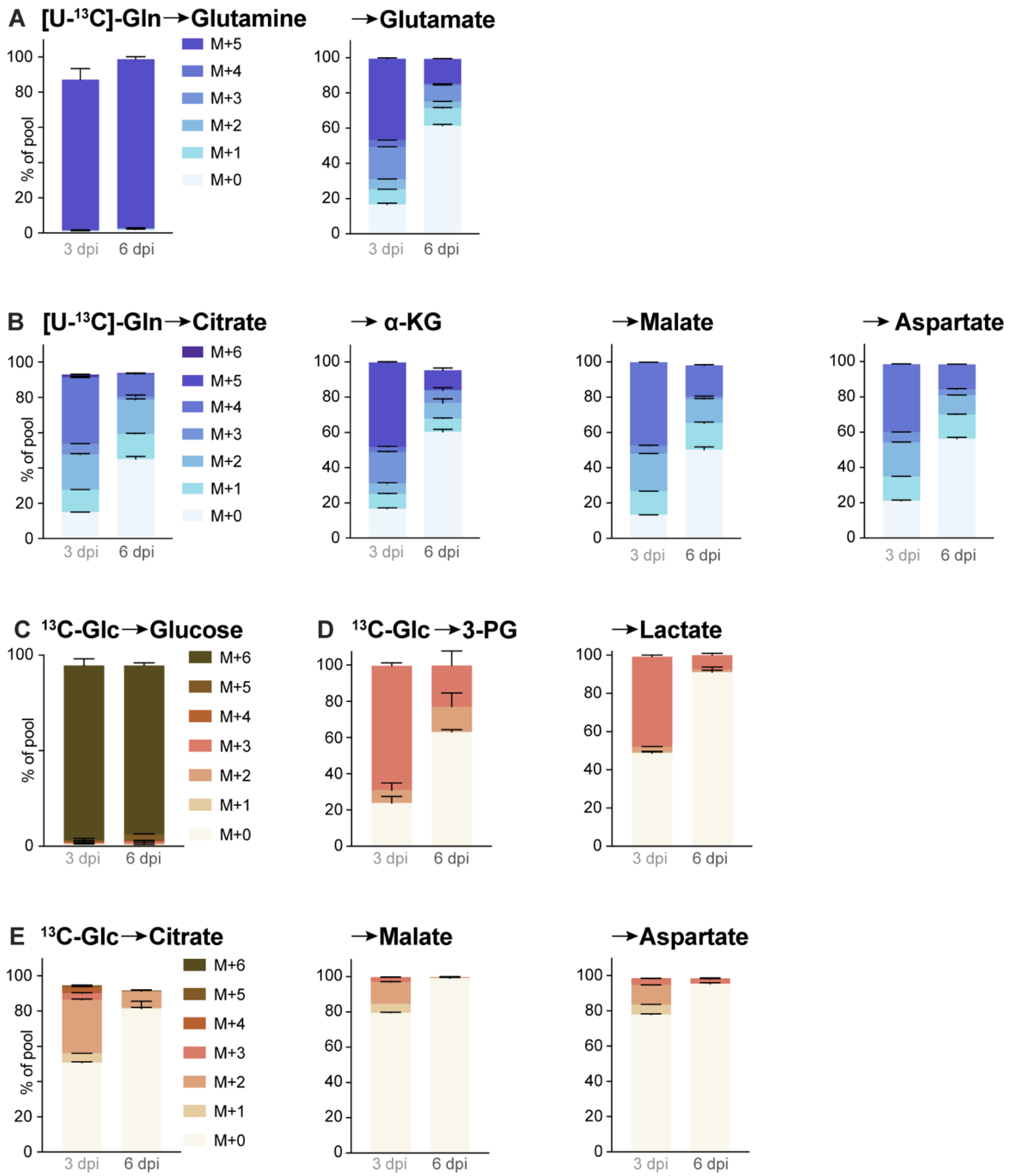


**Fig S5. U-[$^{13}$C]glutamine and U-[$^{13}$C]glucose labeling patterns in T cell metabolites following infusion at different stages of infection. (A-B)** MID for U-[$^{13}$C]glutamine labeling in intracellular **(A)** glutamine and glutamate and **(B)** citrate, α-ketoglutarate, malate, and aspartate in CD8$^{+}$ Teff cells isolated from *Lm*-OVA-infected mice that received a 2h [U-$^{13}$C]-Gln infusion at 3 or 6 dpi. **(C-E)** MID for U-[$^{13}$C]glucose labeling in intracellular metabolites in CD8$^{+}$ Teff cells isolated from *Lm*-OVA-infected mice that received a 2h [U-$^{13}$C]-Glc infusion at 3 or 6 dpi. Shown are the MID for **(C)** glucose, **(D)** glycolytic intermediates 3-phosphoglycerate (3-PG) and lactate, and **(E)** TCA cycle-derived metabolites (citrate, malate, aspartate). Data represent the mean ± SEM for biological replicates (n = 5-6).

**Fig. S6**

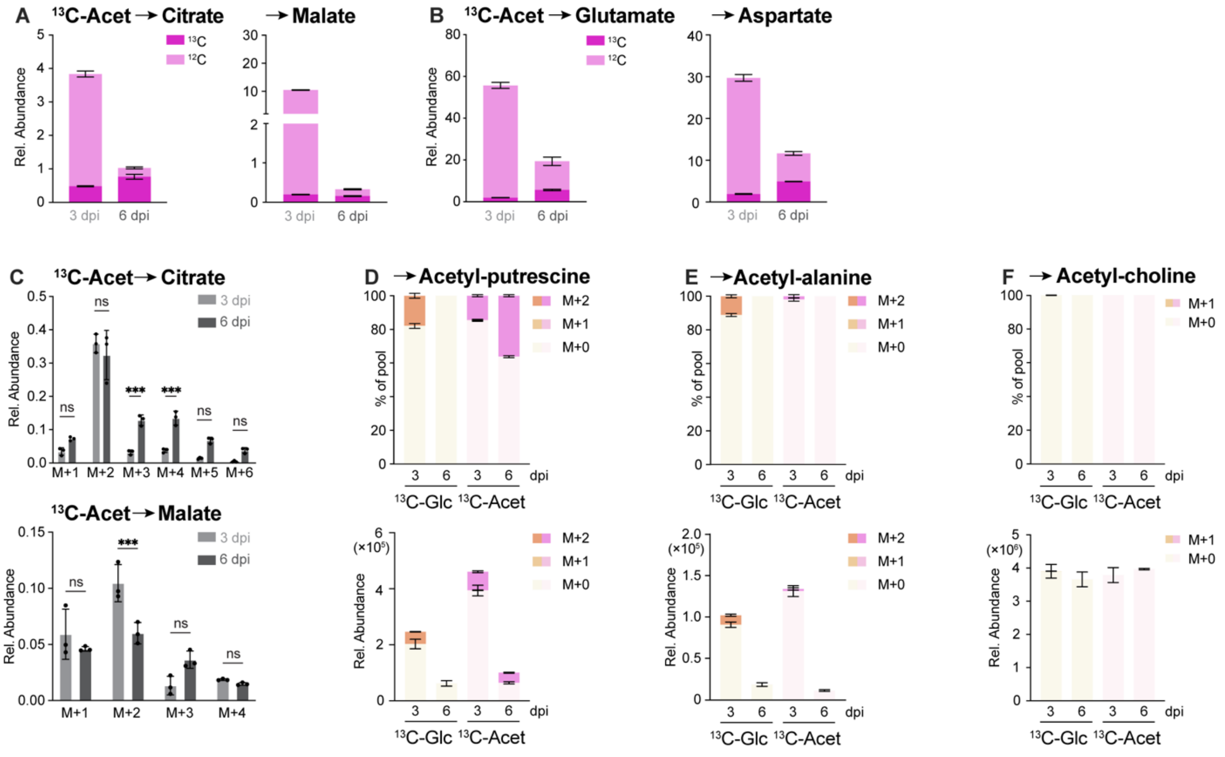


**Fig. S6. U-[$^{13}$C]acetate labeling patterns in T cell metabolites following infusion at different stages of infection.** **(A-C)** U-[$^{13}$C]acetate labeling patterns in intracellular metabolites in CD8$^+$ OT-I T cells isolated from *Lm*-OVA-infected mice at 3 and 6 dpi. Animals received 2h infusion with U-[$^{13}$C]acetate prior to cell isolation and metabolite extraction. Relative abundance of $^{12}$C- versus $^{13}$C-labeled intracellular **(A)** TCA cycle intermediates (citrate, malate) and **(B)** amino acids (glutamate, aspartate). **(C)** Relative abundance of U-[$^{13}$C]acetate-labeled isotopologues of citrate (top) and malate (bottom). **(D-F)** U-[$^{13}$C]acetate and U-[$^{13}$C]glucose labeling patterns in acetylated metabolites in CD8$^+$ OT-I T cells isolated from *Lm*-OVA-infected mice at 3 and 6 dpi as in **(A)**. Animals received 2h infusion with U-[$^{13}$C]acetate or U-[$^{13}$C]glucose prior to cell isolation and metabolite extraction. Shown are the % labelling (top) and relative abundance (bottom) for intracellular **(D)** acetyl-putrescine, **(E)** acetyl-alanine, and **(F)** acetyl-choline. Data represent the mean ± SEM for biological replicates (n=3-6).
